# Deriving Schwann Cells from hPSCs Enables Disease Modeling and Drug Discovery for Diabetic Peripheral Neuropathy

**DOI:** 10.1101/2022.08.16.504209

**Authors:** Homa Majd, Sadaf Amin, Zaniar Ghazizadeh, Andrius Cesiulis, Edgardo Arroyo, Karen Lankford, Sina Farahvashi, Angeline K. Chemel, Mesomachukwu Okoye, Megan D. Scantlen, Jason Tchieu, Elizabeth L. Calder, Valerie Le Rouzic, Abolfazl Arab, Hani Goodarzi, Gavril Pasternak, Jeffery D. Kocsis, Shuibing Chen, Lorenz Studer, Faranak Fattahi

**Author notes:** Correspondence: Lorenz Studer, The Center for Stem Cell Biology Developmental Biology Program Memorial Sloan-Kettering Cancer Center 1275 York Ave, Box 256, New York, NY 10065, Faranak Fattahi, Department of Cellular and Molecular Pharmacology, Eli and Edythe Broad Center of Regeneration Medicine and Stem Cell Research University of California, San Francisco, 600 16th St., GH S572F, San Francisco, CA 94158.

## Abstract

Schwann cells (SCs) are the myelinating and non-myelinating glia of the peripheral nervous system (PNS) and are essential for its function. Defects in SCs are associated with many PNS disorders including diabetic peripheral neuropathy (DPN), a condition affecting millions of patients. Here we present a strategy for deriving and purifying SCs from human pluripotent stem cells (hPSCs). The scalable cultures of SCs allow basic and translational applications such as high-resolution molecular and functional characterization, developmental studies, modeling and mechanistic understanding of SC diseases and drug discovery. Our hPSC-derived SCs recapitulate the key molecular features of primary SCs and are capable of engrafting efficiently and producing myelin in injured sciatic nerves in rats. We further established an hPSC-based *in vitro* model of DPN that revealed the selective vulnerability of human SCs to hyperglycemia-induced cytotoxicity. We established a high-throughput screening system that identified a candidate drug that counteracts glucose-mediated cytotoxicity in SCs and normalizes glucose-induced transcriptional and metabolic abnormalities in SCs. Treatment of hyperglycemic mice with this drug candidate rescues sensory function, prevents SC death, and counteracts myelin damage in sciatic nerves suggesting considerable potential as a novel treatment for DPN.

## INTRODUCTION

Schwann cells (SCs) are vital components and the major glial cells of the peripheral nervous system (PNS). They are crucial for the development, structural maintenance and function of the nerves and exhibit a remarkable ability to promote neural repair following injury (Jessen and Mirsky, 2005; Lavdas et al., 2008). SCs support axons by forming insulating myelin sheaths and Remak bundles, and provide essential neurotrophic factors. Schwann cells develop from the neural crest (NC) via a Schwann cell precursor (SCP) intermediate that is highly proliferative and migratory. SCPs further differentiate into immature SCs that ultimately give rise to mature myelinating or non-myelinating SCs. In addition to SCs, SCPs can give rise to other derivatives (SCPDs) such as melanocytes (Adameyko et al., 2009; Bonnamour et al., 2021; Nitzan et al., 2013). SC defects are involved in genetic and acquired PNS disorders such as Charcot-Marie-Tooth disease, Schwannomatosis, Guillain-Barre Syndrome and diabetic peripheral neuropathy (DPN) for which there are currently no faithful disease models or effective therapies.

Understanding the development and function of SCs and their roles in PNS health and disease has broad basic and translational implications. However, access to authentic models of human SCs at large scale has been a major challenge. Here, we developed hPSC differentiation strategies for efficient derivation of SCs that recapitulate molecular features and function of primary SCs. We characterized the diversity of cell types in our cultures using a combination of imaging and high-resolution transcriptomic profiling and identified novel markers and molecular signatures for human SC subtypes. We further validated the engraftment potential of these cells upon transplantation into a rat model of peripheral neuropathy. Finally, we leveraged hPSC-derived SCs to model the most common cause of peripheral neuropathy, i.e. diabetic peripheral neuropathy (DPN) that affects 30-60% of diabetic patients (Callaghan et al., 2012) and is the leading cause of diabetes-related hospital admissions and nontraumatic lower-extremity amputations (Boulton et al., 2005). The pathogenesis of DPN is complex involving vascular disease, hyperglycemia, hypoxia and oxidative stress that result in cytotoxicity and progressive degeneration of peripheral nerves (Simmons and Feldman, 2002). While symptoms arise from neuronal dysfunction, it is unclear whether sensory neuron damage is the primary event in DPN, and there is evidence that SC degeneration and peripheral demyelination may be contributing factors (Eckersley, 2002). Dissecting cell type specific mechanisms is challenging using current animal models of DPN given the involvement of various non-cell autonomous factors including systemic vascular abnormalities. We utilized hPSC-derived SCs and sensory neurons to determine cell type specific vulnerabilities to high glucose, establish an alternative human-based model of DPN and identify potential therapeutic candidates.

## RESULTS

### Derivation and prospective isolation of SC lineages from hPSCs

We previously established hPSC differentiation protocols to access various NC lineages including enteric and sensory neurons (Barber et al., 2019; Chambers et al., 2012; Fattahi et al., 2016; Tchieu et al., 2017). However, there are currently no methods for the efficient derivation of authentic Schwann cells from hPSCs. Our past efforts of deriving SCs relied on the prolonged, 2-3 months, culture of NC-enriched progenitor cells to obtain a small proportion of gliogenic cells (Lee et al., 2007). More studies reported on the derivation of SC-like cells from hPSCs but did not show molecular authenticity through gene expression profiling and failed to demonstrate functional myelination (Huang et al., 2017; Kim et al., 2017; Liu et al., 2012; Ziegler et al., 2011). While the mechanisms of SC specification during human development remain to be elucidated, SCs are thought to arise from SOX10+ NC cells in a stepwise process. Based on studies in the mouse and chick embryos, NC first gives rise to SC precursors that are competent to associate with neuronal fiber bundles in the developing nerves. The associated neurons produce NRG1 which promotes the survival and further differentiation of SC precursors (SCPs) by activating ERBB3 receptors (Newbern and Birchmeier, 2010). By E13.5 of mouse development, SC precursors give rise to immature SCs which express lineage-specific markers such as GFAP, S100 and POU3F1 while maintaining the expression of SOX10. Terminal differentiation of SCs into myelinating and non-myelinating fates continues for extended time periods and concludes only after birth (Jessen et al., 2015).

Initial hPSC-based NC differentiation protocols relied on the delamination of putative NC cells from neuroepithelial lineages combined with the prospective isolation of p75+ and/or HNK1+ NC precursors (Bajpai et al., 2010; Lee et al., 2007). While those protocols yield various NC-derived lineages, the levels of SOX10 expression are generally low. In contrast, more directed NC induction protocols based on timed exposure to activators of WNT signaling show robust induction of SOX10 in the majority of cells by day 11 of differentiation (Barber et al., 2019; Fattahi et al., 2016; Menendez et al., 2011; Mica et al., 2013; Tchieu et al., 2017). Upon further culture, those hPSC-derived NC cells can be directed into SOX10+ melanocytes (Mica et al., 2013) but also give rise to SOX10-mesenchymal and neuronal precursors (Fattahi et al., 2016; Lee et al., 2007; Mica et al., 2013; Tchieu et al., 2017). Since SOX10 is a key marker in the SC lineage (Finzsch et al., 2010), we first screened for conditions that maintained its expression in cultured NC precursors. We determined the percentage of SOX10+ cells in 2D or 3D NC cultures in the presence of modulators of EGF, FGF, WNT, Notch, TGFβ, BMP and endothelin 3 signaling. We observed that activation of WNT signaling through CHIR99021 exposure and treatment with FGF2 resulted in the maintenance of SOX10 expression in 3D aggregates that we refer to as developing precursors (**Figure 1 A and B**).

**Figure 1:**
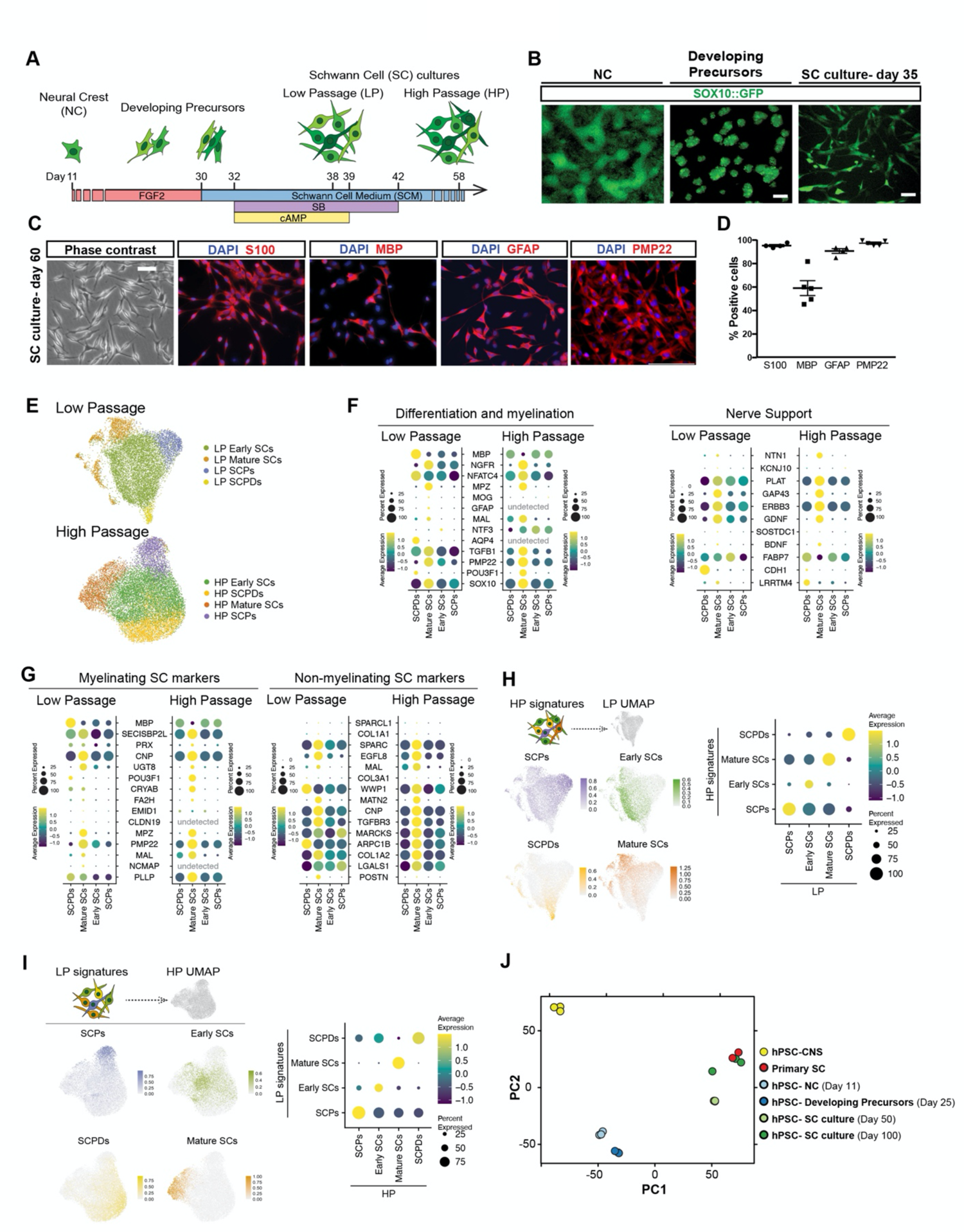
Deriving Schwann cells from hPSCs. **A)** Schematic illustration of the protocol for deriving developing precursors and Schwann cell (SC) cultures from hPSCs-derived neural crests (NCs). **B)** *SOX10::GFP* expression at day 11, 25 and 35 of differentiation. Scale bars = 100 µm in B left and middle panel and 25 µm in B right panel. **C)** Representative immunofluorescence images of hPSC-derived SCs for Schwann lineage markers at day 60. Scale bar= 25 µm. **D)** Quantification of markers in (**D**). **E)** UMAP visualization of scRNA-seq data for low passage (Day 38) and high passage (Day 58) SC cultures. SCPD: Schwann cell precursor derived; SCP: Schwann cell precursor; SC: Schwann cells. **F)** Dot plot of the scaled average expression of SC differentiation and myelination (left) and nerve support (right) markers in single cell RNA-seq data of low and high passage Schwann cells. **G)** Dot plot of the scaled average expression of top 15 primary mouse myelinating (mySC) and non-myelinating (nmSC) Schwann cell markers in single cell RNA-seq data of low and high passage Schwann cell culture. **H)** Module scoring of top 100 high passage (HP) Schwann cell type specific DE marker genes in low passage (LP) Schwann cell types (left). Feature (left) and dot plot (right) visualizations are depicted. **I)** Module scoring top 100 low passage (LP) Schwann cell type specific DE marker genes in high passage (HP) Schwann cell types (left). Feature (left) and dot plot (right) visualizations are depicted. **J)** Principal component analysis (PCA) of NC cells, developing precursors, human primary Schwann cells, and hPSC-derived SC cultures at day 50 and day 100 of differentiation in comparison with central nervous system (CNS) precursors.

Further treatment of developing precursors with Schwann cell media (SCM) containing FGF2, SB431542 and dbcAMP enabled the induction of additional SC markers such as *GFAP*, *POU3F1*, *PMP22*, *MBP*, *AQP4*, and *MPZ* and upregulation of genes involved in glial-neuronal interactions and support including *GDNF*, *ERBB3*, and *GAP43* among others (**Figure S1 A and B**). These cultures can be passaged and maintained for several weeks while retaining the expression of key SC markers S100, MBP, GFAP and PMP22 (**Figure 1 C and D**).

In order to determine the cellular diversity of our hPSC-derived SC cultures, we performed single cell RNA-sequencing (scRNA-seq) at two differentiation time points; low passage (LP, day 38) and high passage (HP, day 58). Unbiased clustering of both LP and HP datasets revealed four transcriptionally distinct cell types namely SCPs, early SCs, mature SCs and SCP derivatives (SCPDs) (**Figure 1E and Figure S2A**). These cell types differentially expressed canonical markers of SC differentiation and function (**Figure 1F**). For example, mature SCs in both LP and HP cultures expressed higher levels of myelinating and non-myelinating SC markers such as *PMP22* and *NGFR*. Nerve support markers and neurotrophic factors including *ERBB3*, *GDNF NGF*, *BDNF* and *GAP43* were also enriched in mature SCs particularly in the HP cultures (**Figure 1F and Figure S2B**). We module scored the LP and HP cell types for a number of SC functional gene sets including neurotrophic factors, neurotransmitter receptors and transcription factors (**Table S1**) and detected differential expression of many neurotransmitter receptors and postsynaptic signal transmission genes by our Schwann cell types (**Figure S2C**). We also identified transcription factors that were specifically expressed by each population in LP and HP cultures. For example, early SCs were enriched for E2F7 and E2F8 while mature SCs expressed SOX10 and FOXO1, POU3F2, TBX19 more specifically. POU6F2 was the common enriched transcription factor in LP and HP SCPDs (**Figure S2D**).

To validate the authenticity of our hPSC-derived SCs, we assessed the expression of top 15 differentially expressed myelinating and non-myelinating Schwan cell specific genes derived from a primary mouse single cell transcriptomics dataset previously published by Segal and colleagues (Tasdemir-Yilmaz et al., 2021) (**Figure 1G, Table S2**). We detected the expression of these markers in both our LP and HP cultures with cell type specific expression patterns (**Figure 1G**). Some markers such as MPZ and MATN2 were specifically expressed by a single cluster namely mature SCs. However, the majority of other genes showed differential transcript levels between cell types while not exclusively expressed in a single population (**Figure 1G**). Interestingly, our LP and HP mature SC populations were highly enriched for both myelinating and non-myelinating markers indicating that our hPSC-derived SCs reliably express markers of authentic SCs.

The proliferative capacity of our SC cultures enables their expansion and scalability. To characterize the proliferation potential of our cell types, we determined the proportional distribution of cell cycle phases within individual LP and HP populations (**Figure S3 A and B**). As cultures transition from low to high passage, all cell types progressively exit the cell cycle. This is particularly evident in SCPs and early SCs that are predominantly cycling in low passage (**Figure S3 A and B**). This is in agreement with the slower proliferation rate of our cultures as they age (not shown).

To assess whether our cultures retain their identity after long-term expansion, we evaluated the lineage relationship between the LP and HP cell types by module scoring the transcriptional signature of LP cell types in HP cell types and vice versa (**Figure 1 H and I**). Notably, each cell type signature was most similar to its corresponding cell type in the other dataset (**Figure 1 H and I**). The similarity between corresponding LP and HP clusters was further demonstrated when we performed clustering on a merged dataset and obtained the same cell populations (**Figure S3 C and D**). Bulk transcriptomics analysis of cultures at different time points demonstrated that hPSC-derived developing precursors were closely related to early NC cells while SC cultures, particularly in higher passages, showed a gene expression pattern closely matching primary adult human SCs (**Figure 1J**).

To characterize the diversity in these cultures in higher resolution, we performed further sub-clustering and revealed two early SCs and two SCPDs populations (**Figure S3E**). While early SCs 1 and 2 separated solely based on their cell cycle phase distributions (**Figure S3F**), SCPDs subclusters were predominantly LP or HP specific (**Figure S3G**). To determine if LP and HP SCPDs were functionally distinct, we performed gene ontology (GO) enrichment analysis of their top 250 differentially expressed genes. Interestingly, despite high level expression of canonical melanocytic genes such as MITF, MLANA and PMEL, LP SCPDs were enriched for myelin production terms, such as cholesterol and lipid metabolism pointing to a dual melanocyte-SC identity (**Figure S2A and Figure S3H**). On the other hand, HP SCPDs displayed enrichment for melanin synthesis and pigmentation (**Figure S3H**) indicating that as cultures age SCPDs become more melanocytic.

Strategies for prospective isolation enables the generation of pure and high quality SC populations from heterogeneous cultures. To enable fluorescence-activated cell sorting (FACS) based purification of hPSC-SCs, we screened a library of 242 antibodies for human surface antigens that specifically mark GFAP+ SCs (**Figure S4 A and B**). We identified 11 surface antibodies that stained >20% of the GFAP+ SCs of which CD44, CD49e, CD81 and CD98 labeled most of the target population (**Figure S4B**). Analysis of surface marker expression in our LP and HP scRNA-seq datasets revealed transcripts that were specifically enriched in each cell type (**Figure S4C**). Interestingly, this list included 9 of the 11 surface marker hits identified by the antibody screening. Among these, CD46, CD146, CD147 and CD166 were enriched in mature SCs in both datasets while CD9, CD49e and CD171 were enriched only in HP mature SCs. CD44 was highly expressed in SCPDs in both LP and HP cultures while CD81 enrichment was specific to the LP population (**Figure S4C**). Further validations revealed that CD98 was the only marker specifically expressed in SCs but not in NCs or SCPs (**Figure S4D**). These populations expressed CD49D, a marker previously shown to label early SOX10+ NC lineages (Fattahi et al., 2016). Collectively these data demonstrate that our hPSC differentiation system generates scalable and proliferative human SC cultures that can be further enriched using FACS.

### hPSC-SCs promote neuronal maturation and myelination *in vitro* and engraft into injured sciatic nerves in rats

SCs play fundamental roles in maintaining and protecting the structure and function of the peripheral nerves. Myelinating SCs are specialized glial cells that form lipid rich myelin sheaths around the axons and enable fast neuronal signal propagation in the PNS. Since our hPSC-derived mature SCs express genes involved in myelination and lipid metabolism in high levels (**Figure 1 F and G, Figure S3H**), we set out to identify and characterize myelinating SCs (mySCs) in our LP and HP mature SCs. We module scored a curated list of top differentially expressed primary mouse mySCs and canonical myelinating genes (Calder et al., 2015) in our LP and HP mature SCs (**Figure 2A, Table S3**). We identified >25% of mature SCs in each dataset as myelinating (**Figure 2A**). LP and HP mySCs were specifically enriched for a number of neurotrophic factors and neurotransmitter and postsynaptic transmission genes (**Figure S5 A and B**). For example, members of TGF and FGF protein families were highly enriched in mySCs compared to other mature SCs in both datasets (**Figure S5A**). BDNF expression was specific to HP mySCs (**Figure S5A**). To define the unique functional properties of these cells we performed pathway enrichment analysis using GO BP, KEGG and Reactome gene sets on the significantly upregulated genes in LP and HP mySCs (**Figure 2B**). Interestingly, of the top 50 significantly enriched pathways, we identified multiple pathways related to axon development, myelination, neuronal development, synapse assembly, cell adhesion and cell motility. These are well established physiological features of myelinating SCs. HP mySCs upregulated pathways related to extracellular matrix organization, cell adhesion and cell motility among others. This is intriguing given the known contribution of SCs in depositing and organizing ECM components, formation of lamellipodia and cytoplasmic protrusions and forming contact and recognizing axons during radial sorting and myelination. These indicate our LP and HP mySCs are equipped with the molecular programs that enable myelinating SCs to perform their function.

**Figure 2:**
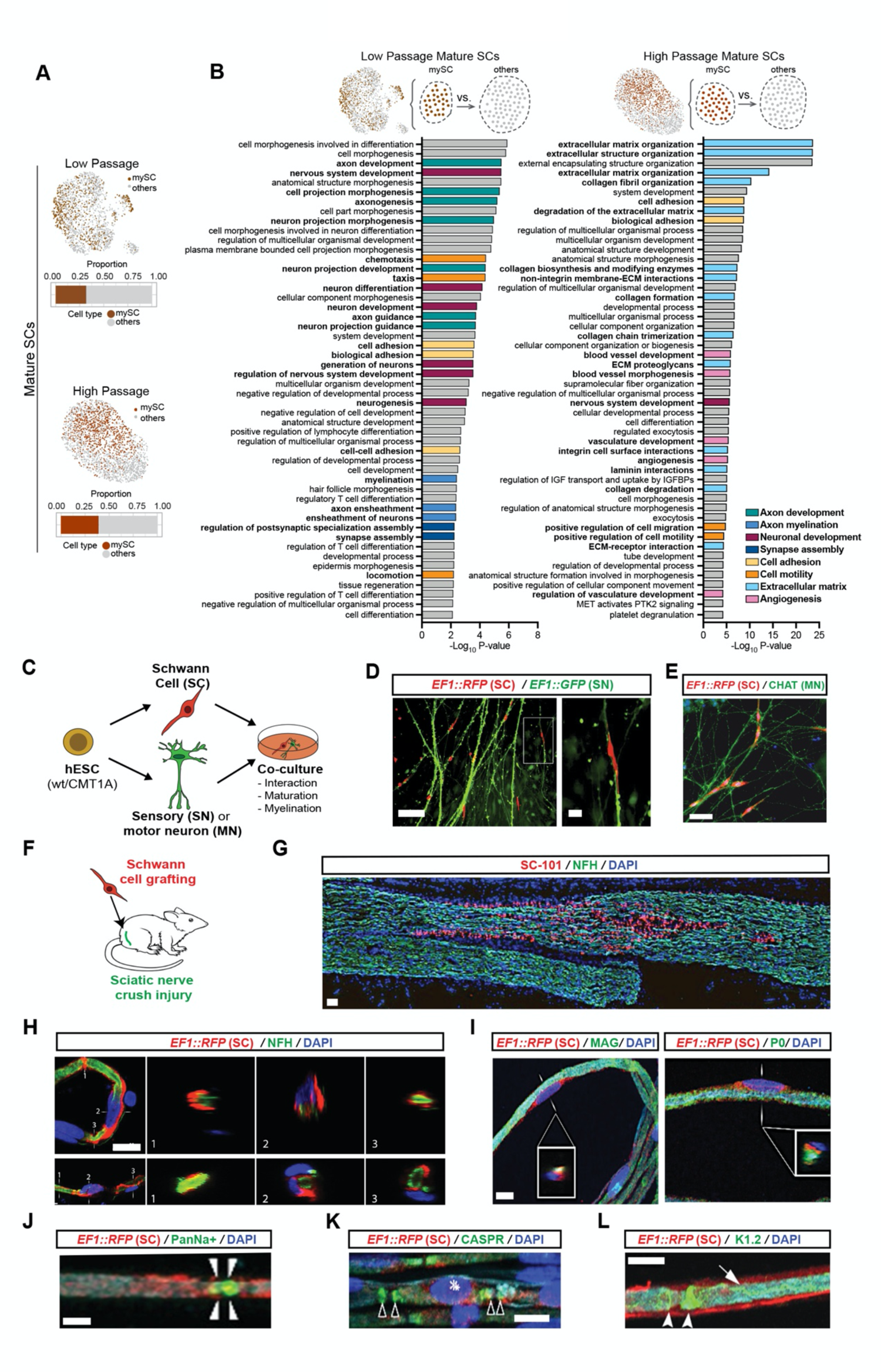
hPSC-derived Schwann cells myelinate hPSC-derived sensory neurons and engraft in injured rat sciatic nerves. **A)** Feature plots of mature SC clusters isolated from low (top) and high (bottom) passage culture single cell RNA-seq data with dark colors indicating SCs identified as myelinating (mySC). Bar plots show the relative population of mySCs. **B)** Pathway enrichment analysis of top 250 DE genes of myelinating mature SCs in low (left) and high (right) passage cells. Top 50 pathways from combined GO BP, Reactome and KEGG analysis are shown. **C)** Schematic illustration of the hPSC-SC co-cultures with hPSC-derived sensory or motor neurons. **D)** Physical association of hPSC-SCs with hPSC-sensory neurons. **E)** Physical association of hPSC-SCs with hPSC-derived motor neurons. **F)** Schematic illustration of hPSC-SC transplantation in adult rat sciatic nerves. RFP+ hPSC-derived Schwann cells were injected following nerve crush at the site of injury (adult Cyclosporin-A treated SD rats). **G)** Immunofluorescence staining of grafted sciatic nerves for human specific nuclear marker SC101 at 8 weeks post transplantation. **H)** Confocal analysis of teased sciatic nerve fibers for RFP (grafted human cells), axonal marker (NFH) and DAPI. **I)** Confocal analysis of teased nerve fibers for RFP (grafted human cells), myelin markers MAG and P0 and DAPI. **J-L**) Confocal analysis of teased nerve fibers for RFP and node markers Pan-Na+ (sodium channel, arrow heads, H), CASPR (arrow heads, **I**) and Kv1.2 (K+ channel, arrow heads, **J**). Scale bars= 100 µm in B left panel, 20 µm in B right panel, 0.2 µm in C, 100 µm in, 20 µm in F and G and 10 µm in H-J.

Since cell adhesion molecules (CAMs) play an important role in SC association with axons, nerve components and the ECM, and cell adhesion was amongst the most significantly enriched GO terms in both our LP and HP mySCs (**Figure 2B**), we sought to determine the specific CAMs that were enriched in mySC populations. We module scored a list of cell adhesion molecules combining cell-cell and cell-matrix gene sets in our LP and HP mature SCs and identified many CAMs that were specifically depleted and enriched in LP and HP mySCs relative to the other mature SCs (**Figure S5C**). For example, HLA-DR that was also a hit in our SC antibody screening (**Figure S4B, Figure S5C**) was enriched in both LP and HP mySCs. The proinflammatory interleukin IL18 was depleted while MAG and KIT were enriched in both mySC populations (**Figure S5C**). Members of the contactin protein family (CNTN1, 4 and 6) that are axon-associated CAMs and play roles in the formation of axon connections in the developing nervous system were specifically enriched in HP mySCs (**Figure S5C**). Similarly, HP mySCs were highly enriched for PLXNB3 which is important for axon guidance and cell migration (**Figure S5C**).

To assess the ability of hPSC-derived SCs to functionally interact with neurons, we established co-cultures with hPSC-derived sensory (Chambers et al., 2012) and motor neurons (Calder and Tchieu, 2015) (**Figure 2C**). RFP-labeled SCs (day 60) were mixed with GFP-labeled sensory neurons (day 50) and analyzed at 72 hours of co-culture. SCs associated closely with sensory neurons by aligning along their processes (**Figure 2D**). Similarly, co-cultures of SCs and hPSC-derived motor neurons (day 25) showed robust interaction along neuronal fibers (**Figure 2E**). The protracted process of human cell maturation in hPSC-derived lineages is a major hurdle in the field. Glial cells such as astrocytes have been shown to promote the functional maturation of hPSC-derived CNS neurons (Tang et al., 2013). To assess the impact of SCs on maturation, we performed calcium imaging in hPSC-derived motoneurons at day 40 and 70 of differentiation (15 and 55 days of co-culture). Interestingly, there was a marked increase in the calcium response of stage-matched motoneurons co-cultured with SCs (**Figure S6A**). By day 70, the responsiveness to glutamate and KCl stimulations further improved and remained distinct from cultures containing motor neurons only (**Figure S6B**). Our findings demonstrate the capability of hPSC-SCs to modulate neuronal function *in vitro*. In order to determine whether hPSC-SCs are functional *in vivo* and are capable of producing myelin, we asked whether they could survive and engraft in a rat model of sciatic nerve injury. We depleted endogenous SCs through a mechanical crush of the nerve and injected RFP-labeled hPSC-SCs at the site of injury (**Figure 2F**). The transplanted SCs could be readily detected at eight weeks after nerve injection using the human-specific nuclear marker SC101 (**Figure 2G**). Transplanted hPSC-SCs were in close contact with the host neurons (**Figure 2H**) and expressed the myelin markers MAG and P0 (**Figure 2I**). In mature myelinated fibers, sodium channels are localized at nodes of Ranvier: the site of action potential electrogenesis. This area is flanked by a CASPR-expressing domain (juxtaparanodal region) where the axon membrane is in close contact to myelin membrane. Adjacent to the CASPR+ region is an axon membrane domain marked by the expression of potassium channels. Remarkably, we observed appropriate localization of both sodium and potassium channels in axons that were wrapped by RFP-labelled hPSC-SCs (**Figure 2J-L**). These studies demonstrate the ability of hPSC-SCs to engraft and produce myelin that is appropriately associated with nerve fibers and the nodes of Ranvier in injured adult peripheral nerves.

These results demonstrate that our hPSC-derived cultures of functional SCs offer a framework for modeling pathologies in which SCs play central roles in disease initiation and progression. For example, a large subset of Charcot-Marie-Tooth (CMT) patients suffer from debilitating myelin defects caused by genetic mutations. Importantly, genes associated with CMT including de-myelinating CMT1, axonal CMT2, and intermediate CMT are expressed by our SC cultures confirming their applicability for modeling CMT pathophysiology in future studies (**Figure S7**).

### hPSC-derived SCs enable modeling, mechanistic understanding and treating diabetic peripheral neuropathy

In addition to rare inherited defects such as CMT, SCs are associated with a broad range of other neuropathies. The most prominent form of an acquired neuropathy is diabetic peripheral neuropathy (DPN) which results from the progressive degeneration of peripheral nerves (Simmons and Feldman, 2002). While symptoms arise from neuronal dysfunction, it is unclear whether sensory neuron damage is the primary event in DPN, and there is evidence that SC degeneration and peripheral demyelination may be contributing factors (Eckersley, 2002). As a proof of concept, we set out to leverage our human hPSC differentiation system to model DPN by investigating the effect of high glucose on sensory neurons and SCs (**Figure 3A**).

**Figure 3:**
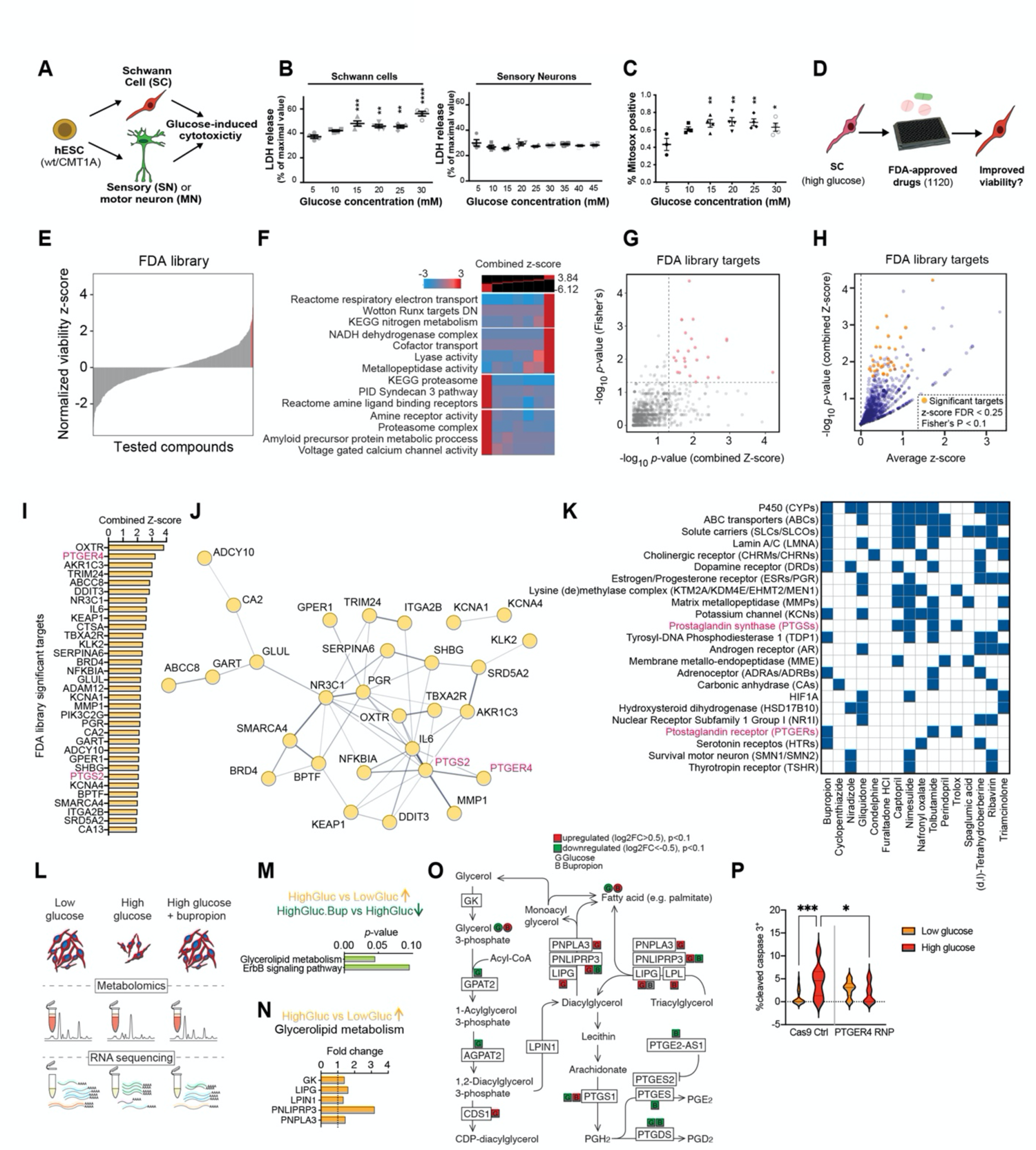
Schwann cells are selectively vulnerable to high glucose exposure. **A)** Schematic illustration of the experimental paradigm for modeling diabetic nerve damage in hPSC-derived cell types. **B)** Lactate dehydrogenase (LDH) cytotoxicity analysis of hPSC-derived SCs and sensory neurons in response to exposure to different glucose concentration using LDH activity assay. **C)** Oxidative stress measurement of hPSC-derived SCs exposed to increasing concentration of glucose. Statistical analysis was performed using one-way ANOVA comparing values to the low glucose (5 mM) condition. ns, not significant, p-values are: * p<0.05; ** p<0.01. **D)** Schematic illustration of high-throughput drug screening for identification of compounds that enhance the viability of high glucose-treated hPSC-SCs. **E)** Presentation of the distribution of library compounds by their corresponding normalized viability z-score. **F)** Gene-set enrichment analysis using iPAGE for the library compounds targets identifies GO terms associated with hits improving and worsening SC viability. **G)** p-value correlation plot to identify the genes that are most likely the targets of the effective treatment. Normalized z-score from all the treatments associated with a gene are integrated. In addition, to assess the enrichment of individual genes among those that are targets of the treatments with increased z-scores, a Fisher’s exact test is performed. The plot shows the correlation between the p-values. **H)** One-sided volcano plot showing the average *z*-score vs. -log of *p*-value for all genes with positive *z*-scores. The genes that pass the statistical thresholds of combined *z*-score FDR<0.25 and Fisher’s *p*-value <0.1 are marked in gold. **I)** Identified target genes (marked gold in H) ranked by their combined z-score. **J)** Protein-protein interaction network of the identified target genes (listed in I) constructed by STRING database. Minimum required interaction score was set to 0.4 and edge thickness indicates the degree of data support. **K)** Predicted targets of hits in (K) compiled from the following databases: BindingDB (Liu et al., 2007), Carlsbad (Mathias et al., 2013), Dinies (Yamanishi et al., 2014), PubChem bioassays (Kim et al., 2019), SEA (Keiser et al., 2007), Superdrug 2 (Siramshetty et al., 2018) and SwisTargetPrediction (Gfeller et al., 2014). **L)** Schematic illustration of the unbiased metabolite and transcriptional profiling of differentially treated SCs. **M)** Pathway enrichment analysis of genes upregulated in high glucose and downregulated upon BP treatment. **N)** Glycerolipid metabolism enzymes upregulated in high glucose condition. **O)** Glycerolipid metabolism schematic adopted from KEGG shows changes in the enzymes and metabolites in response to high glucose and BP treatments in SCs. HighGluc: high glucose, LowGluc: low glucose, HighGluc.Bup: high glucose plus bupropion, LowGluc.Bup: low glucose plus bupropion **P)** PTGER4 KO in SCs rescues high glucose treated SCs. CRISPR-Cas9 mediated knocking out of PTGER4 in SCs protects them against increased levels of cleaved caspase 3 (marker of apoptosis) under high glucose condition as measured by flow cytometry. p-values are: * p<0.05; ** p<0.01; *** p<0.001.

Sensory neurons showed no overt toxicity at glucose levels of up to 45 mM. In contrast, hPSC-derived SC cultures were exquisitely sensitive to even moderately increased glucose levels (**Figure 3B**). Treatment with high glucose induced oxidative stress in SC cultures as measured by MitoSOX staining (**Figure 3C**).

Given the exquisite glucotoxicity in SCs, strategies that prevent glucose-mediated cellular damage in SCs may represent novel therapeutic opportunities for treating DPN. We established a high-throughput screening (HTS) assay to measure the viability of hPSC-SCs in the presence of 30 mM glucose and the Prestwick library containing 1,120 small molecules of approved drugs (FDA, EMA or other regulatory agencies) (**Figure 3D, Figure S8A**). We identified several hit compounds that significantly increased SC viability under high glucose conditions (**Figure 3E, Table S4**). Gaining mechanistic insights on the protective effect of these hits could shed light on the mechanism of glucotoxicity in SCs. Given that the library compounds target many different cellular pathways, we sought to determine the shared pathways among candidate drugs that improved SC viability under the high glucose condition using our previously established analysis approach (Samuel et al., 2020).

We first predicted drug protein interactions for the entire Prestwick library using the similarity ensemble approach (SEA)(Keiser et al., 2007). We then added normalized z-score across all compounds that target each protein to calculate weighted combined z-score for each protein and used it for iPAGE GO analysis (Goodarzi et al., 2009). Among the GO terms associated with the SC protecting drug candidates we identified oxidative phosphorylation (OXPHOS), nitrogen metabolism and metallopeptidases (**Figure 3F**). We validated the expression of these candidate pathways in our HP cultures using module scoring analysis (**Figure S8B**). To determine the degree to which positive z-scores were enriched among the drugs targeting each protein, we performed a Fisher’s exact test. Through this analysis, we identified 33 proteins as significant drug targets filtered based on average combined z-score>0, false discovery rate (FDR)<0.25 and Fisher’s p<0.1 (**Figure 3 G-I**).

We performed a protein-protein interaction network analysis to identify interactions between our 33 significant drug targets using the STRING database (Szklarczyk et al., 2019). In the resulting network, IL6, NR3C1, PGR and PTGS2 had the highest degree centrality (**Figure 3J**). In addition, for a more comprehensive target prediction, we generated a list of potential targets for the top hits derived from the HTS dataset (**Figure S8C**) or computational predictions that use network-based and similarity-based algorithms (**Figure 3K, Table S5**). Intriguingly, many of the potential target proteins were shared between multiple hits; including potassium channels (KCNs), estrogen and progesterone receptors (ESRs and PGRs), prostaglandin synthases (PTGSs) and prostaglandin receptors (PTGERs). Notably, the predicted target pattern of tolbutamide resembled that of our top hit, bupropion (BP) (**Figure 3K**).

Our top hit compound, BP, is a widely used antidepressant marketed as Wellbutrin^®^. BP showed a dose-dependent effect on rescuing the viability of high glucose treated SC cultures (**Figure S8D**). In line with previous reports on glucose-mediated activation of oxidative stress response (Corrêa-Silva et al., 2018; Kowluru et al., 2003), we observed the activation of cellular inflammatory response via NF-kB p65 localization to the nucleus in SCs exposed to high glucose, a phenotype that was counteracted by BP treatment (**Figure S8E**).

To understand the mechanism of glucotoxicity in SC cultures and its rescue with BP, we performed unbiased transcriptional and metabolite profiling (**Figure 3L**). We compared gene expression profiles of SCs treated with low glucose and high glucose with or without BP treatment followed by pathway enrichment analysis of differentially expressed genes (**Figure S9 A and B**). Of the 1,559 SC transcripts that were enriched or depleted in response to high glucose, 66 showed reversed expression pattern in cells treated with BP (**Figure S9A**). Of these, *PTGER4* was the only gene in common with the list of predicted targets of BP (**Figure 3 I and K, Table S5**). The affected processes in cells exposed to high glucose included several metabolic and cell cycle signaling pathways and BP treatment modulated pathways related to DNA replication and transcription as well as cell cycle and stress response (**Figure S9B**).

To better understand the metabolic consequences of high glucose and BP treatments, we performed metabolomics and integrated the data with our bulk RNA sequencing results (**Figure S10 A and B, Figure S11A**). The detected primary metabolites fell into seven categories based on their pattern of abundance in response to glucose and BP (**Figure S10 A and B**). Many of the metabolites were either accumulated or depleted in SCs exposed to high glucose. BP treatment led to a reversed response in a subset of these metabolites in particular groups 4 and 7. Metabolic pathway enrichment analysis suggested modulations in citric acid cycle, urea cycle, amino acid metabolism, glycolysis and gluconeogenesis (**Figure S10 A and B**). In parallel, BP treatment led to an increase in the detected levels of TCA cycle metabolites succinate, fumarate, citrate, α-ketoglutarate and malate (**Figure S10, Figure S11A**). The concentration of citrate was reduced in response to high glucose and was reversed by BP treatment. Citrate transporter *SLC13A2* expression followed the same trend (**Figure S10, Figure S11A**). Exposure to high glucose resulted in elevation of cellular urea that was accompanied by increased transcript levels of the plasma membrane urea transporter *SLC14A2* (**Figure S10, Figure S11A**). Both urea concentration and its transporter mRNA level were decreased in the presence of BP (**Figure S10, Figure S11A**). We observed an increase in cellular pyruvate in response to high glucose that was lowered by BP treatment (**Figure S10, Figure S11A**). Cellular lactate level was reduced after BP treatment which was accompanied by downregulation of membrane monocarboxylate transporters *SLC16A3, SLC16A14, SLC5A2* RNA levels (**Figure S10, Figure S11A**). It has been previously reported that in some cell types elevated glucose levels can activate the polyol pathway (Oates, 2002). The polyol pathway metabolizes excess intracellular glucose into sorbitol and subsequently into fructose via two enzymatic steps catalyzed by aldose reductase (AR) and sorbitol dehydrogenase (SDH), respectively. Osmotic and oxidative stress triggered by the polyol pathway have been proposed as mediators of tissue damage in response to high glucose based on studies in the lens more than 50 years ago (Van Heyningen, 1959). Sorbitol accumulation has been implicated in peripheral nerve damage in multiple animal models of diabetes (Mizisin, 2014; Oates, 2002). We asked whether hPSC-SCs show increased sorbitol levels in response to high glucose as a potential mechanism of their selective vulnerability. In agreement with studies in the mouse (Maekawa et al., 2001; Mizisin and Powell, 1993), we observed a higher AR to SDH ratio in hPSC-derived SCs compared to sensory neurons (**Figure S11B**). Furthermore, SCs but not sensory neurons showed increased levels of sorbitol when exposed to high glucose (**Figure S11C**). BP treatment reduced sorbitol accumulation in SCs in a dose-dependent manner (**Figure S11D**). Similarly, we observed an increase in the cellular level of fructose in SCs treated with high glucose and BP treatment countered this effect (**Figure S11E**). This is in agreement with the protective effect of aldose reductase inhibitors in diabetic SCs (Hao et al., 2015). Collectively, these results point to a global metabolic shift in high glucose treated SCs that is reversed by BP treatment.

Pathway enrichment analysis of transcripts that were upregulated in response to high glucose and reversed in response to BP revealed glycerolipid metabolism and ErbB signaling pathways (**Figure 3 M**). Transcriptional changes in the glycerolipid metabolism pathway (**Figure 3N**) were accompanied by corresponding changes in pathway metabolites measured in the metabolomics dataset (**Figure 3O**). Increased tri-and diacylglycerol degradation into free fatty acid and glycerol was suggested by upregulation of *LIPG* and *PNLIPRP3* transcripts in response to high glucose treatment (**Figure 3N**). This is interesting given the crucial role of lipid metabolism in myelin production and SC physiology (**Figure S3H**). Further, through the action of prostaglandin synthases, the glycerolipid metabolic pathway is directly linked to the prostaglandin metabolism. This piqued our interest for multiple additional reasons. For example, prostaglandin E2 receptor (PTGER4) and prostaglandin-endoperoxide synthase 2 (PTGS2) were among the top significant protein targets identified by our high-throughput drug screening (**Figure 3I**). Remarkably, they were both part of the protein-protein interaction network with PTGS2 showing a high degree network centrality (**Figure 3J**) and PTGSs and PTGERs were part of the protein families shared between our top drug hits (**Figure 3K**). Moreover, PTGER4 transcript had a reversed pattern of expression in response to high glucose in presence and absence of BP (**Figure S9A**). We aimed to investigate the effect of PTGER4 in SC glucotoxicity using a genetic approach. We knocked out PTGER4 in the Schwann cells using CRISPR-Cas9 ribonucleoproteins (RNP) and observed a significant reduction in cleaved caspase-3 in response to high glucose treated cells indicating a lower sensitivity to glucotoxicity in the absence of PTGER4 (**Figure 3P**).

### Bupropion rescues the disease phenotypes in a mouse model of DPN

Given the remarkable ability of BP in rescuing the viability of hPSC-SCs *in vitro*, we next assessed the therapeutic potential of BP in a mouse model of DPN. We treated wild type C57BL6 mice with the pancreatic beta cell specific toxin streptozotocin (STZ) which leads to beta cell death, impaired insulin production and hyperglycemia in mice (Wu and Huan, 2001). In DPN, sensory nerve damage commonly leads to loss of sensation in the extremities. We evaluated the impact of BP treatment in STZ-treated mice by measuring thermal sensation as a readout of sensory nerve function and by histological analysis of the sciatic nerve to assess structural damage (**Figure 4A**). STZ-treated mice showed a dramatic increase in blood glucose levels independent of BP treatment as compared to non-diabetic control animals (**Figure S12A**) indicating that BP treatment does not affect glucose levels. Hyperglycemic mice maintained without BP treatment showed a delayed response to thermal stimulation at seven and eight weeks post-STZ treatment. Remarkably, BP treated diabetic mice showed no significant difference in response time compared to normal, non-diabetic animals (**Figure 4B**). Histological analysis revealed a marked increase in the percentage of TUNEL+ apoptotic cells in the sciatic nerves of STZ mice. BP+STZ treated animals showed significantly fewer apoptotic cells than animals with vehicle+STZ treatment (**Figure 4 C and D**). Finally, we evaluated the impact of STZ and BP treatment on peripheral myelin using transmission electron microscopy. We observed a large percentage of fibers with damaged myelin in the sciatic nerves of STZ treated animals which was significantly reduced in BP+STZ treated animals (**Figure 4 E and F**). These data indicate that BP can partially prevent DPN in STZ treated mice. To assess whether BP is capable of reversing sensory defects and has therapeutic potential in more advanced disease states, we started BP treatment 8 weeks post-STZ in a separate cohort of mice. These mice already showed a delayed response time to thermal stimulation before treatment with BP but showed no significant difference after 4-8 weeks of BP treatment compared to normal mice (**Figure S12B**). These studies demonstrate robust therapeutic effects of BP in the STZ-model of DPN. Collectively these results highlight the promise of hPSC-derived cultures for modeling SC disorders, understanding disease mechanisms and identifying potential therapeutic strategies.

**Figure 4:**
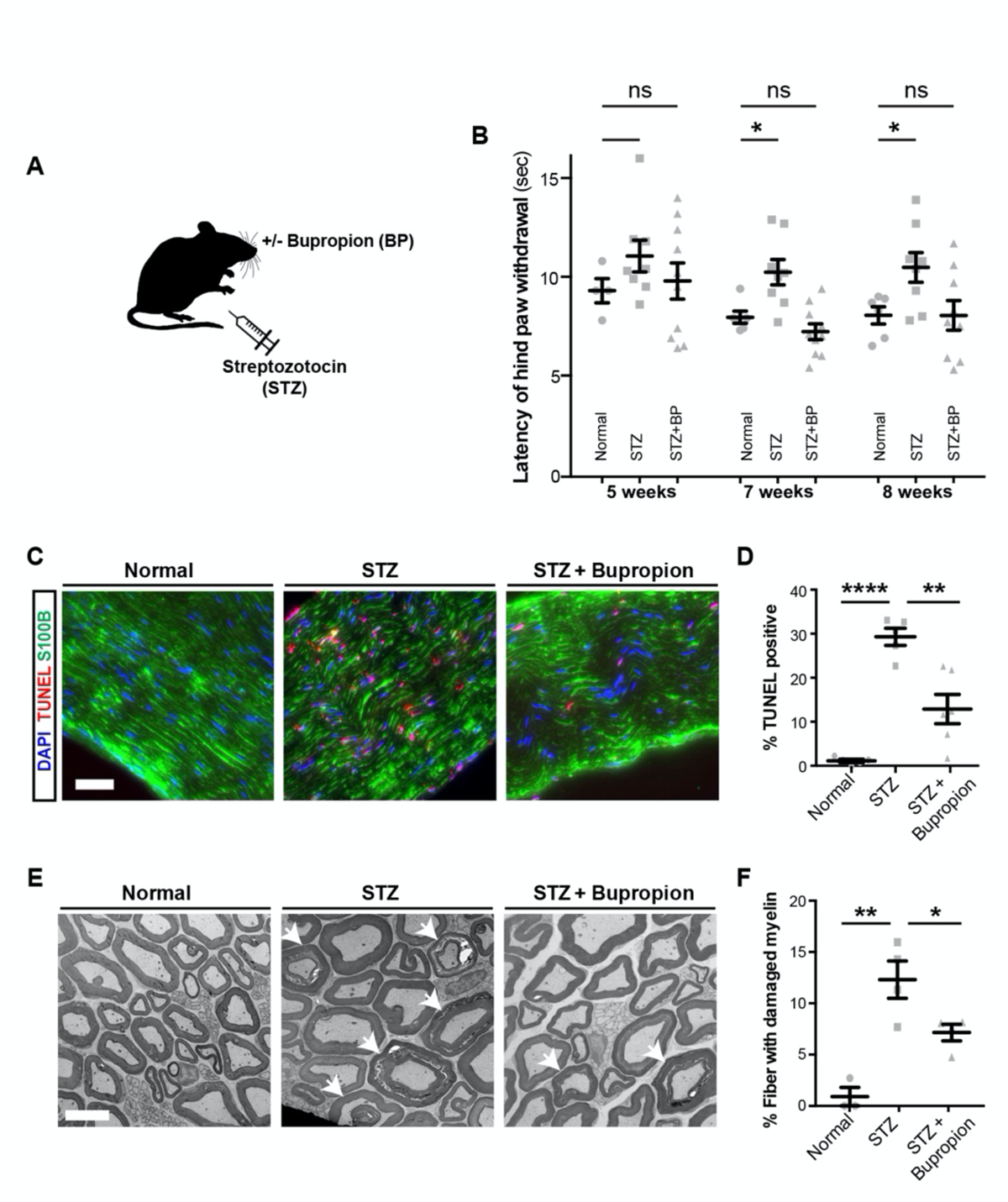
Bupropion treatment prevents diabetic nerve damage in mice. **A)** Schematic illustration of modeling diabetes and Bupropion treatment in mice. **B**) Thermal sensitivity test measuring the latency of hind paw withdrawal in normal mice and mice treated with STZ and Bupropion. **C, D**) TUNEL staining and quantification in sciatic nerves of normal mice and mice treated with STZ and Bupropion. **E, F**) Electron transmission microscopy and quantification of damaged myelin structures in sciatic nerves of normal mice and mice treated with STZ and Bupropion. p-values are: * p<0.05; ** p<0.01; *** p<0.001; **** p < 0.0001. Scale bars= 100 µm in C and 5 µm in E. BP, Bupropion HCL.

## DISCUSSION

Our study reports on the highly efficient derivation of SCs from hPSCs which overcomes the limitations of previous studies such as low yield, protracted differentiation, limited SC maturation and lack of myelination data (Huang et al., 2017; Kim et al., 2017; Lee et al., 2007; Liu et al., 2012, 2012; Ziegler et al., 2011). An important feature of our hPSC-based model is the scalability and purity of the resulting SCs and the ability to culture cells for extended periods without losing SC properties. In contrast, primary SCs tend to rapidly lose their properties upon extended culture which results in the increasing contamination of SC cultures with fibroblast-like cells. Important developmental questions that are now accessible using this novel differentiation technology include the mechanisms controlling the transition from a multipotent NC stem cell to committed SCs and the study of human SC plasticity given data in the mouse suggesting that both melanocytes and parasympathetic neurons can be derived from early SC-lineages (Adameyko et al., 2009; Bonnamour et al., 2021; Espinosa-Medina et al., 2014; Nitzan et al., 2013). Our stepwise SC differentiation protocol generates SCs following the developmental processes thought to occur *in vivo*. It hence provides a powerful framework for lineage tracing studies that would shed light on how NC and SCP fate specification mechanisms lead to the emergence of SCs and other SCPDs such as melanocytes.

Our high-resolution transcriptomics profiling of hPSC-derived SCs is the first study that provides a comprehensive molecular characterization of human SCs at two differentiation stages. In addition to confirming the authenticity of the differentiated Schwann cell types, it allows us to identify specific transcription factors, SC functional markers and surface molecules. Combining these data with high-throughput antibody screening, we identify novel markers for prospective isolation of mature SCs. The identification of CD98 as a surface marker for the prospective isolation of committed SCs represents a powerful tool for such studies. Based on the proof-of-concept data presented here, studies on SC-mediated neuronal maturation and myelination should be other areas of focus. The modeling of PNS pathologies and the development of a drug screening platform for compounds modulating peripheral myelination could be of particular interest. A surprising feature of the cultured hPSC-derived SC is their gene expression pattern that not only confirms SC identity but suggests that pluripotent-derived cells match the expression pattern of adult SC. This is in contrast to most other *in vitro* derived hPSC-lineages such as neurons expressing fetal stage markers (Studer et al., 2015). Autologous SCs are currently being tested for applications in regenerative medicine targeting both PNS and CNS disorders (Lavdas et al., 2008; Rodrigues et al., 2012). Our transplantation data demonstrate robust engraftment of hPSC-SCs in a model of traumatic nerve injury. While future studies are required to assess long-term engraftment and therapeutic potential in models of PNS and CNS injury, our results set a solid foundation for the application of hPSC-SCs in regenerative medicine including spinal cord injury.

We present an hPSC-based DPN model that revealed a selective vulnerability of SCs to diabetes-associated hyperglycemia. While most of our data were obtained in response to high glucose exposure, we observed decreases in SC viability even at more moderate levels of increased glucose. Through our high-throughput screening, we identified candidate drugs that counteract the glucotoxicity in SCs. Interestingly there are many shared proteins between the predicted targets of these drugs. For validation studies, we focused on our top candidate bupropion. Transcriptional and metabolic profiling of stressed and rescued SCs gave interesting insight into the cellular consequences of SC glucotoxicity and the mechanism of BP rescue. By comparing the list of BP predicted targets and the 66 transcripts that had reversed pattern of expression under glucose and BP treatment, we identified PTGER4 as a potential target that mediates SC glucotoxicity. These proteins convert arachidonate to PGH2 that is the precursor for other prostaglandins including PGD2 and PGF2 and PGE2, the latter being the ligand for PTGER4. In addition, PTGER4 is among the significant targets of our HTS library compound analysis of hits that improve SC viability under high glucose condition. Moreover, prostaglandin analogs have been shown some evidence of moderate efficacy in a randomized clinical trial for DPN (Boulton et al., 2005).

The ability of BP to counteract glucose-mediated SC toxicity correlated with a decrease in intracellular glucose and sucrose levels. BP treatment resulted in reduced cellular lactate concentration and its membrane transporters that might be potentially important given the importance of lactate as a fuel in neuron-SC metabolic coupling (Babetto et al., 2020; Domènech-Estévez et al., 2015). Our transcriptional and metabolomics profiling provide evidence on upregulation of glycolysis and downregulation of mitochondrial respiration in SCs in response to high glucose that were rescued by BP treatment. iPAGE gene set enrichment analysis also identified oxidative phosphorylation as pathways by which HTS hits may mediate their protective effects. Interestingly, BP appears to be the only antidepressant drug commonly associated with moderate weight loss in patients rather than a weight gain (Arterburn et al., 2016). It is tempting to speculate whether BP mediated changes in glucose metabolism could be related to those systemic effects. By performing pathway enrichment analysis, we identified significant changes in glycerolipid metabolism under high glucose condition that was reversed upon BP treatment. This is of significant functional relevance considering the role of lipid metabolism and myelin synthesis in SC function.

Importantly, our *in vivo* studies demonstrate that BP treatment can rescue DPN-related behavioral deficits and nerve damage. The fact that BP is a widely used drug should greatly facilitate any future testing in DPN patients. Interestingly, BP has shown some benefit in the treatment of patients suffering from neuropathic pain (Semenchuk et al., 2001) raising the question whether those effects of the drug for alternative indications could also be mediated by its effect on SC vulnerability. In addition to BP, we identified several additional compounds capable or rescuing Schwann cell vulnerability. It will be interesting to determine whether those compounds act via a common or distinct mechanism and whether they show *in vivo* activity in STZ mice comparable to BP.

In conclusion, our findings facilitate human SC-based studies for applications in regenerative medicine and human disease modeling. The work further implicates SC defects in the pathogenesis of DPN and presents BP as an FDA-approved drug that can treat DPN-related damage *in vitro* and *in vivo*.

## ACKNOWLEDGEMENTS

We are very grateful for technical support provided by Harold S. Ralph (Weill Cornell Cell Screening Core) and Leona Cohen-Gould (Weill Cornell histology and electron microscopy Core). The work was supported by grants from UCSF Program for Breakthrough Biomedical Research and Sandler Foundation, the NIH Director’s New Innovator Award (DP2NS116769) and the National Institute of Diabetes and Digestive and Kidney Diseases (R01DK121169) to F.F., the Starr Foundation and by NYSTEM contract C026446 to L.S. and supported by the NIH Cancer Center support grant P30 CA008748; by TRI-SCI 2014-030 to L.S., and S.C., and by the New York Stem Cell Foundation (R-103), NIDDK (DP2 DK098093-01) to S.C. S.C. is a New York Stem Cell Foundation – Robertson Investigator. H.G. is supported by NIH (R01CA240984). H.M. is supported by Larry L. Hillblom Foundation postdoctoral fellowship.

## AUTHOR CONTRIBUTIONS

H.M.: Design of experiments, writing of manuscript, maintenance, directed differentiation of human ESCs, differentiation protocol optimization, protocol gene knockout experiments, biochemical assays, quantitative PCR analyses, metabolomics and bulk RNA-seq data analysis, protein-protein and protein-drug interaction analyses. S.A.: Calcium imaging and in vivo drug treatment assays. Z.G.: Small molecule screening, establishing diabetic mouse model and *in vivo* drug treatment assays. A.C. scRNA-seq and snRNA-seq data analysis, writing of manuscript. E.A. and K.L.: Sciatic nerve transplantation and histological analysis. S.F.: Immunofluorescence imaging and analysis. A.K.C.: Maintenance, directed differentiation of human ESCs, flow cytometry analysis. M.O.: Maintenance, directed differentiation of human ESCs, immunofluorescence imaging. M.D.S.: Maintenance, flow cytometry analysis. J.T.: Bulk RNA-seq data analysis. E.C.: Derivation of motor neurons from hPSCs. V.L.R.: Design and execution of thermal sensitivity test in mice. A.A.: iPAGE analysis. H.G.: Bulk RNA-seq data analysis. G.P.: Study design for mouse behavioral test. J.K.: Design and execution of sciatic nerve transplantation studies. S.C.: Design and interpretation of small molecule screen and follow up experiment. L.S.: Design and conception of the study, data interpretation, writing of manuscript. F.F.: Design and conception of the study, data interpretation, writing of manuscript, maintenance, directed differentiation of human PSCs, developing differentiation protocol, establishing of co-culture assays, cellular, molecular and biochemical assays, histological analyses, small molecule screen and *in vivo* experiments.

## DECLARATION OF INTERESTS

L.S. is a scientific co-founder and paid consultant of BlueRock Therapeutics, and he is listed as an inventor of several patents owned by MSKCC related to hPSC-differentiation technologies including a patent application with F.F., L.S., S.C., G.Z. as inventors describing the technologies reported in the current manuscript.

---

**Figure S1:**
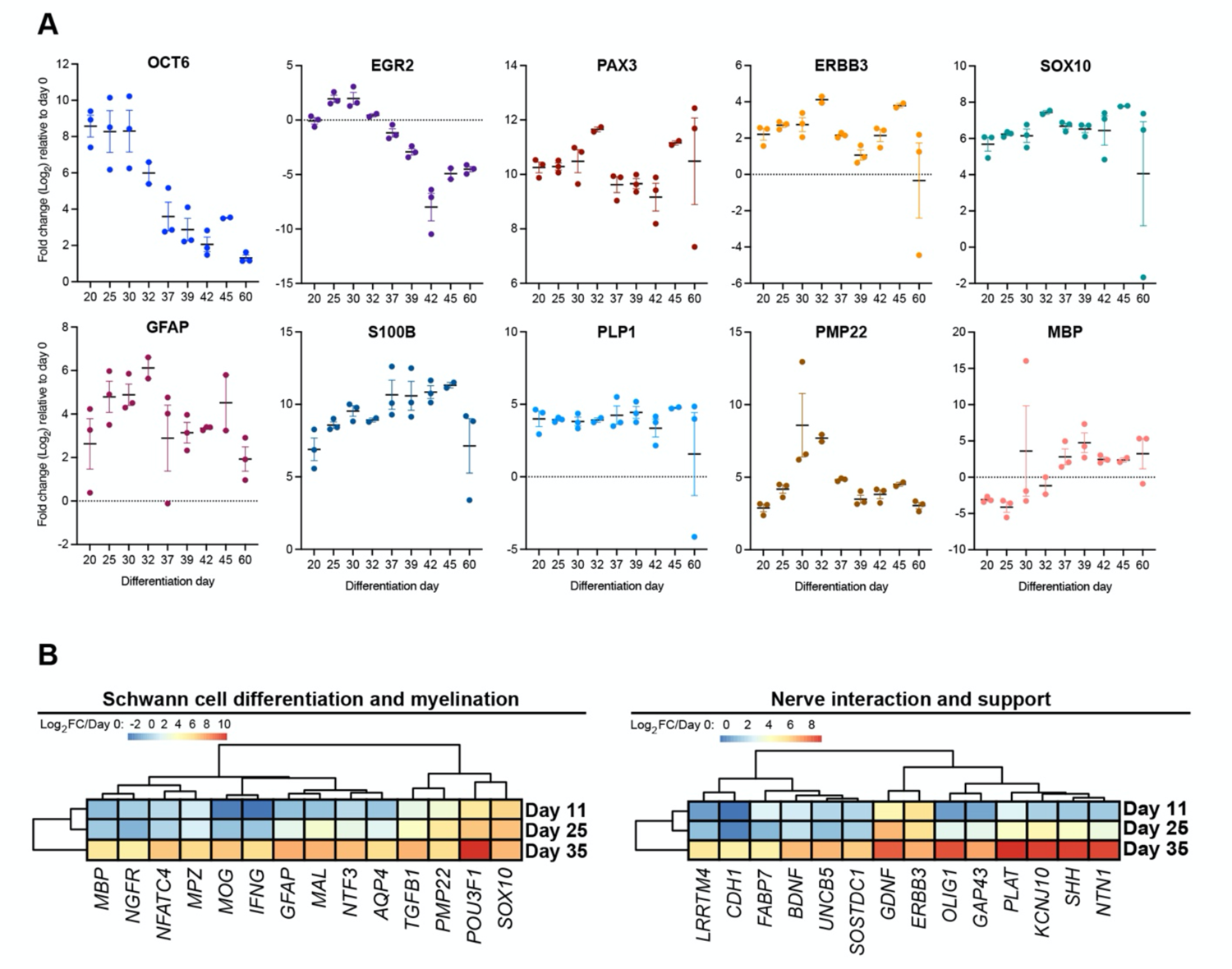
Characterization of hPSC-derived SC lineages. **A)** qRT-PCR for a panel of Schwann cell and precursor markers at different timepoints of the differentiation protocol. Log_2_ fold change relative to D0 of differentiation is shown. **B)** qRT-PCR for a panel of Schwann lineage markers involved in Schwann cell differentiation and myelination (left) and nerve interaction and support (right).

**Figure S2:**
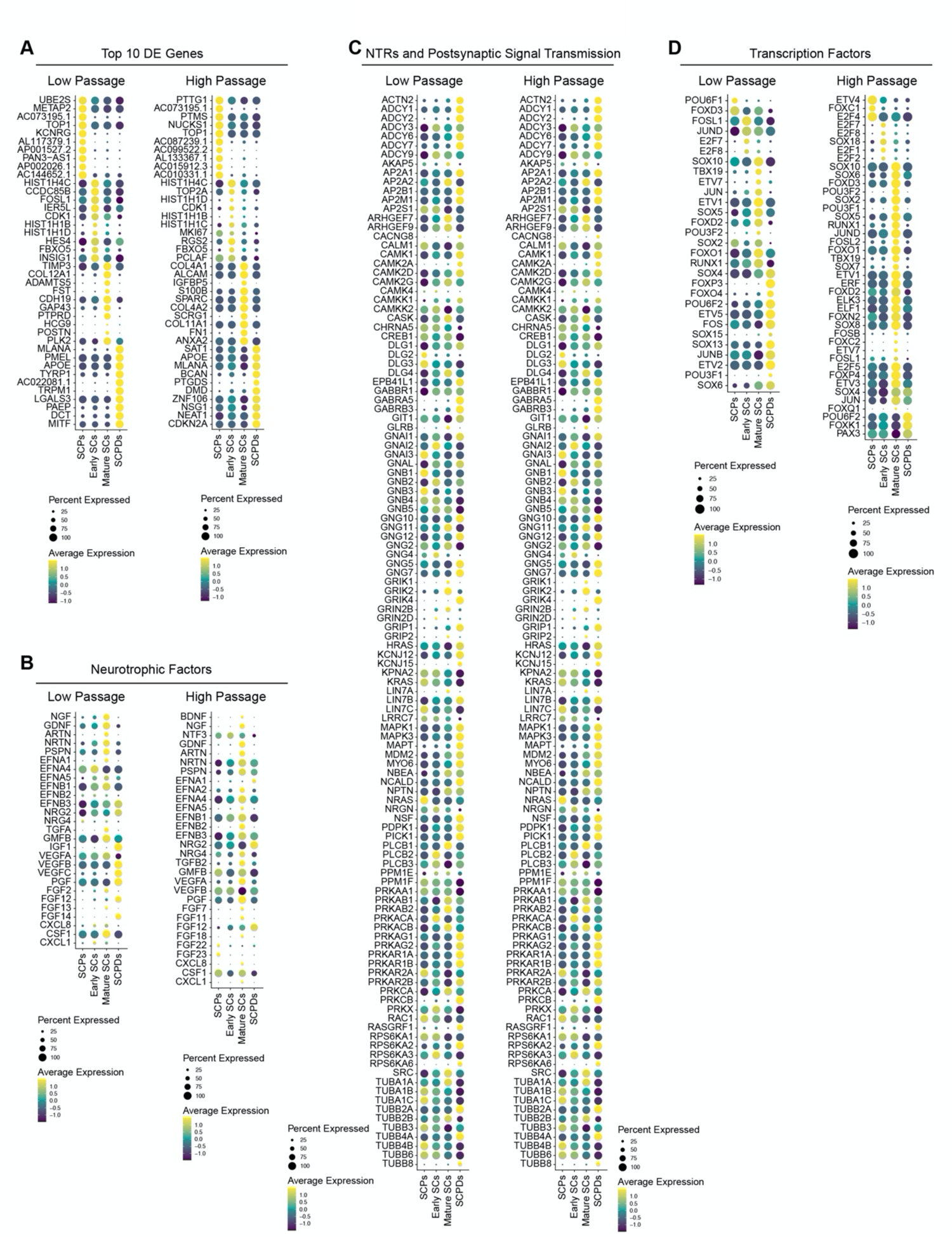
Sub-type specific characterization of hPSC-derived SC lineages. **A**) Dot plot of the scaled average expression of the top ten differentially expressed (DE) genes for each low and high passage Schwann cell types. SCPD: Schwann cell precursor derived; SCP: Schwann cell precursor; SC: Schwann cells. **B-D**) Dot plots of scaled average expression of low and high passage cell type specific neurotrophic factors (**B**), neurotransmitter receptors and postsynaptic signal transmission (**C**), and transcription factors (**D**). All gene sets were prior filtered to include genes expressed in at least 25% of cells in either cell type in low (LP) and high (HP) passage SC culture data.

**Figure S3:**
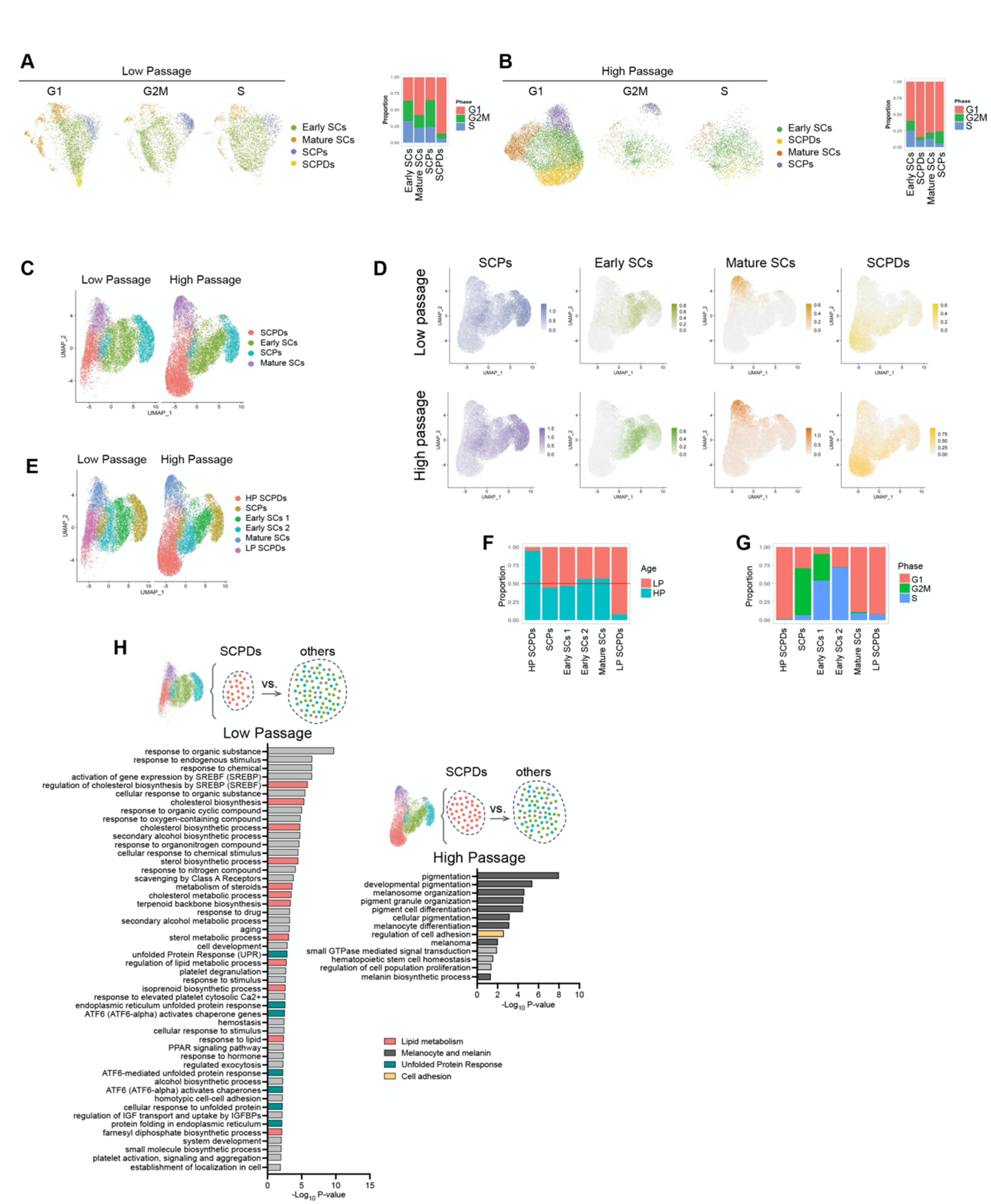
Molecular changes in SC cultures after long-term maintenance. **A and B**) Feature plot (left) and distribution of cell cycle phases (right) in low passage (**A**) and high passage (**B**) Schwann cell cultures **C)** UMAP visualization of merged low and high passage Schwann cell dataset. **D)** Feature plots showing module scoring of top 100 low passage (top) and high passage (bottom) cell type specific DE marker genes in the merged dataset. **E-G**) UMAP visualization, (**E**) distribution of cell cycle phases (**F**), and cell population proportion (**G**) in low and high passage merged dataset depicting subclusters of early SCs and SCPDs. **H**) Gene ontology biological process (GO BP) pathway enrichment analysis on top 250 differentially expressed genes in low and high passage SCPDs.

**Figure S4:**
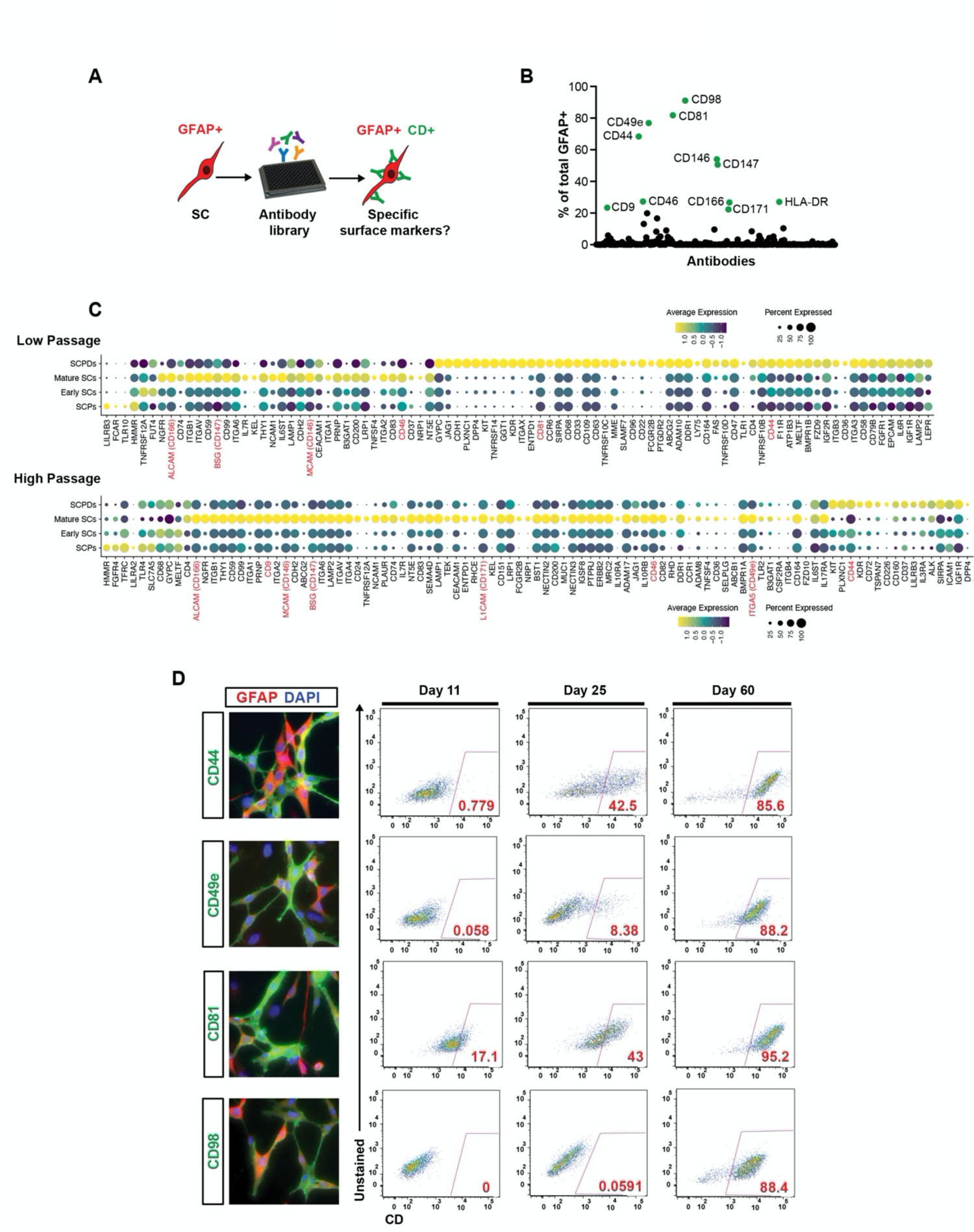
Antibody screen identifies novel surface markers for human SCs. **A)** Schematic illustration of the human antibody screening paradigm. **B)** Primary screening identifies novel surface markers for hPSC-SCs. **C)** Dot plot of the scaled average expression of low and high passage cell type specific surface markers. Surface markers identified in the human antibody screen highlighted in red. Gene set was prior filtered to include genes expressed in at least 25% of cells in either cell type in low (LP) and high (HP) passage SC culture data. **D)** Immunocytochemistry (left), flow cytometry-based (right) validation of surface marker expression at different stages of SC differentiation.

**Figure S5:**
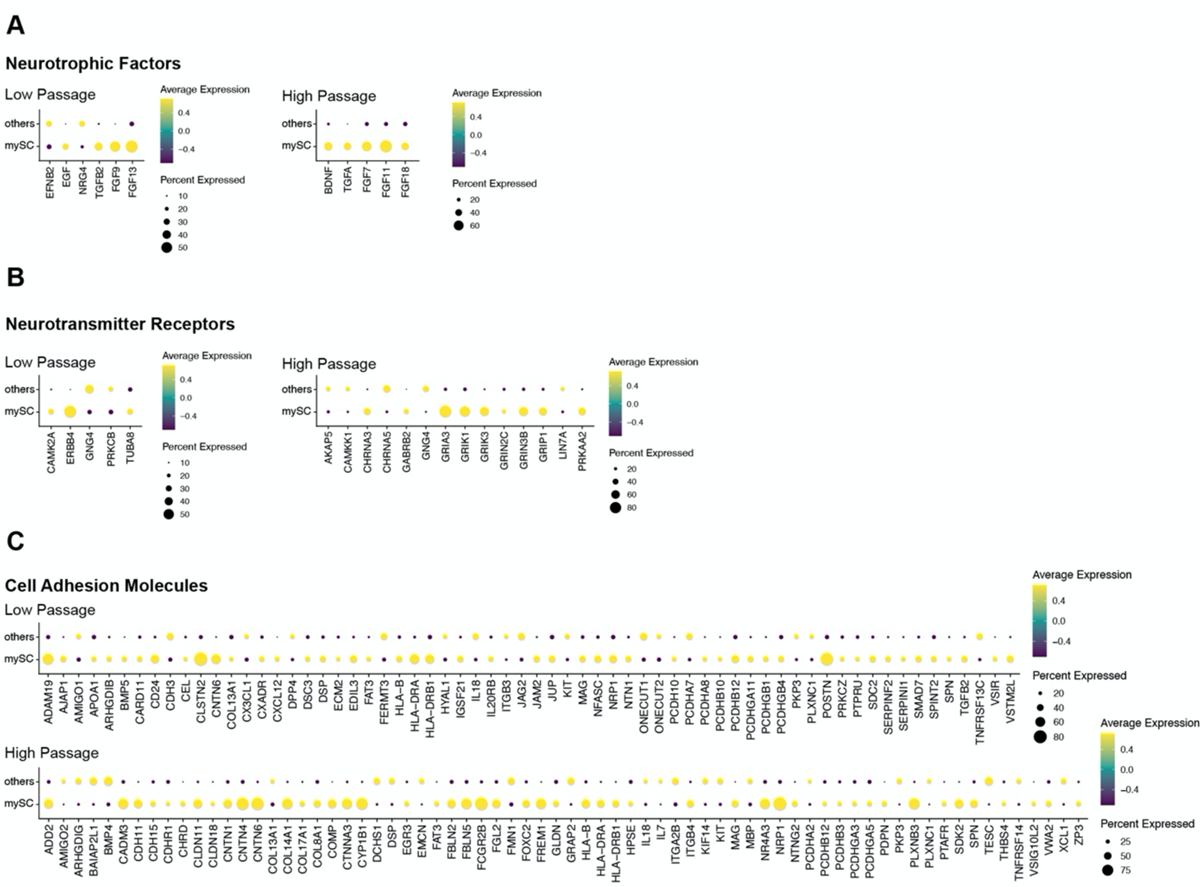
Molecular characterization of myelinating SCs. **A-C**) Dot plots of scaled average expression of specific neurotrophic factors (**A**), neurotransmitter receptors (**B**), and Cell Adhesion Molecules (**C**) in myelinating SCs (myScs) and other cells in low and high passage cultures. All gene sets were prior filtered to include genes expressed in at least 25% of cells in either cell type in low (LP) and high (HP) passage SC culture data.

**Figure S6:**
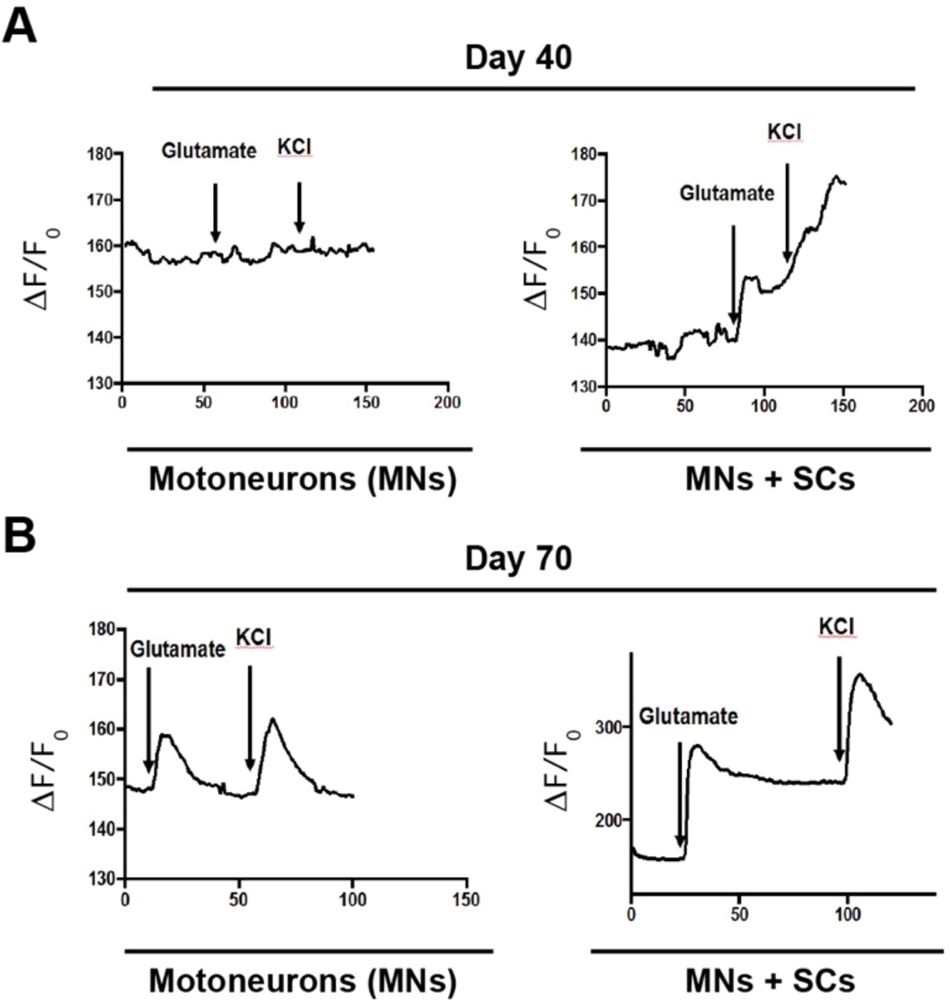
hPSC-SCs accelerate functional maturation of hPSC-motor neurons in co-cultures. **A and B**) Calcium imaging of motor neuron single culture and motor-neuron Schwann co-cultures at day 40 (**A**) and 70 (**B**) of motor neuron differentiation.

**Figure S7:**
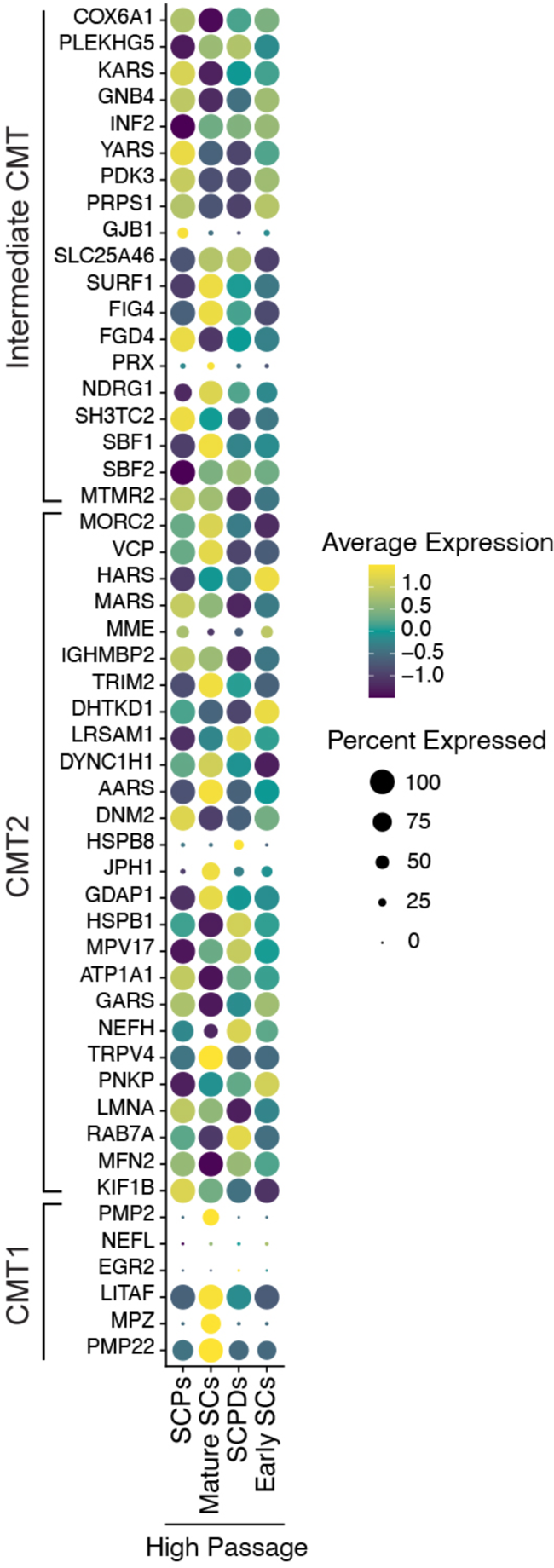
hPSC-SCs express Charcot-Marie Tooth (CMT) disease associated genes. Dot plot of the scaled average expression of Charcot-Marie Tooth (CMT) disease associated genes in high passage cell types

**Figure S8:**
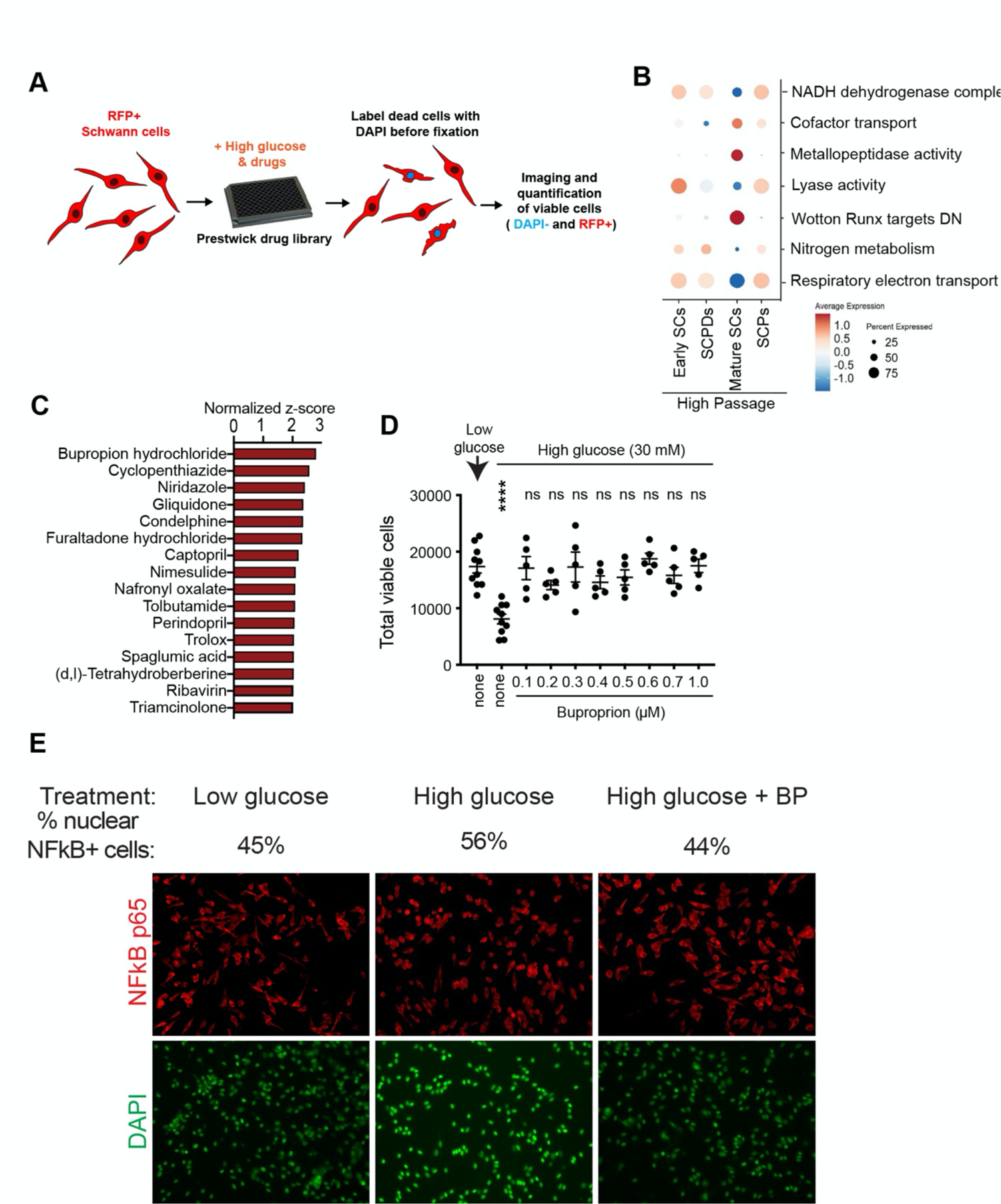
High-throughput screening identifies drugs that rescue glucotoxicity in hPSC-SCs. **A)** Schematic illustration of the high-throughput screening paradigm for identification of drugs that counteract glucotoxicity in hPSC-SCs. **B)** Module scoring of pathways that improve SC viability under hyperglycemia (identified by iPAGE analysis, Figure 3F) in high passage SC dataset. Dot plot (left) and violin plot (right) are shown. **C)** HTS top hits with normalized z-score > 2 (marked red in Figure 3E). **D)** Dose response of BP treatment in hyperglycemic SCs. **D**) Analysis of NF-kB pathway activation by measuring p65 subunit localization to cell nucleus under high glucose. Percentage of nuclear p65 positive SCs in response to high glucose and BP as measured by immunofluorescent imaging are shown. Statistical analysis was performed using one-way ANOVA comparing values to the low glucose (5 mM) condition. ns, not significant, ****, p<0.0001.

**Figure S9:**
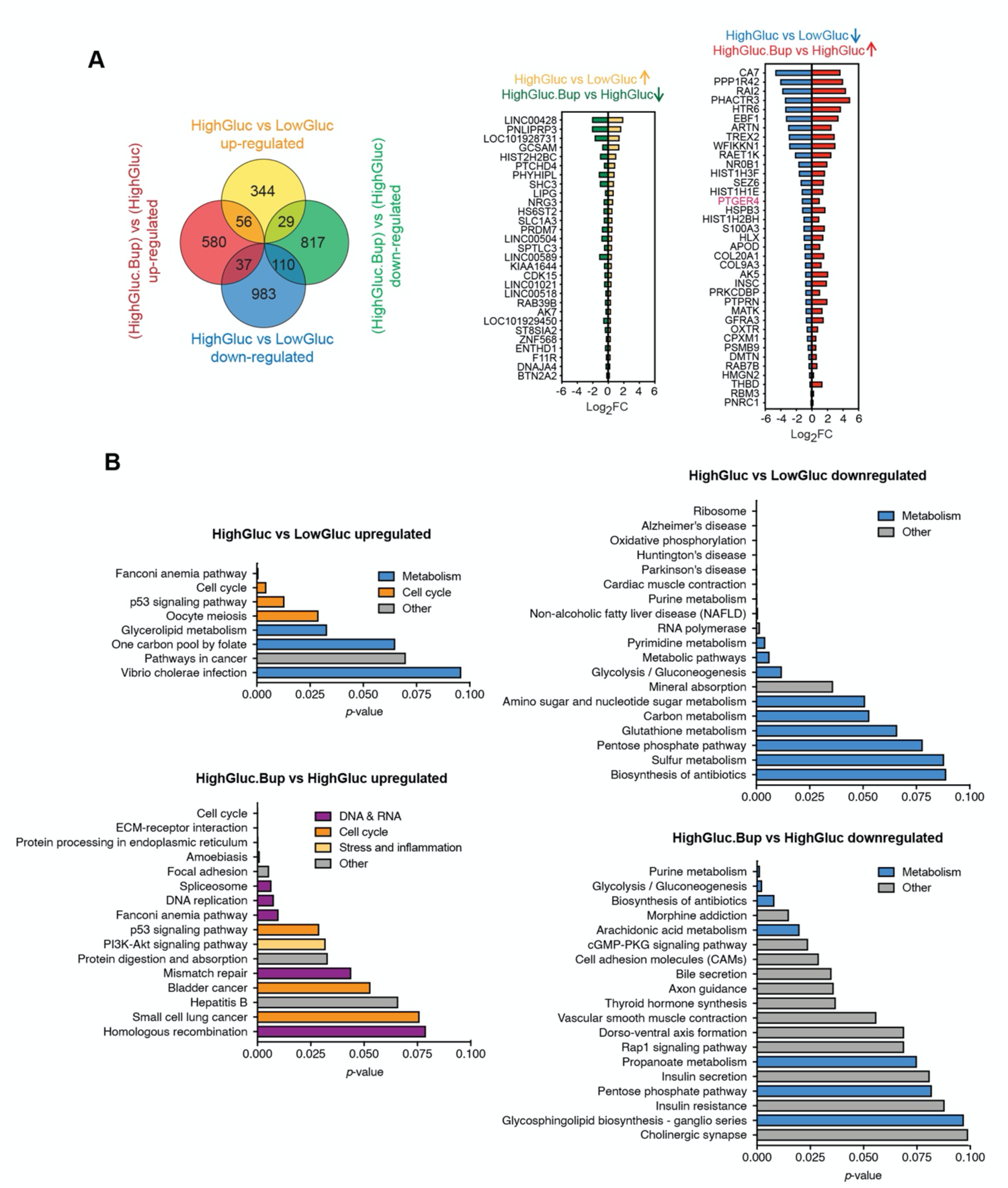
Glucose and BP treatments differentially affect SCs transcriptome. **A)** Venn diagram showing differentially expressed genes in SCs treated with high glucose, plus and minus BP (left). Differentially expressed genes in hyperglycemia that show a reversed pattern of expression when treated with BP (middle and right). **B)** Genes that were differentially expressed when SCs were treated with high glucose, and high glucose plus BP were subjected to pathway enrichment analysis using DAVID (Huang et al., 2009a, 2009b). HighGluc: high glucose, LowGluc: low glucose, HighGluc.Bup: high glucose plus bupropion, LowGluc.Bup: low glucose plus bupropion

**Figure S10:**
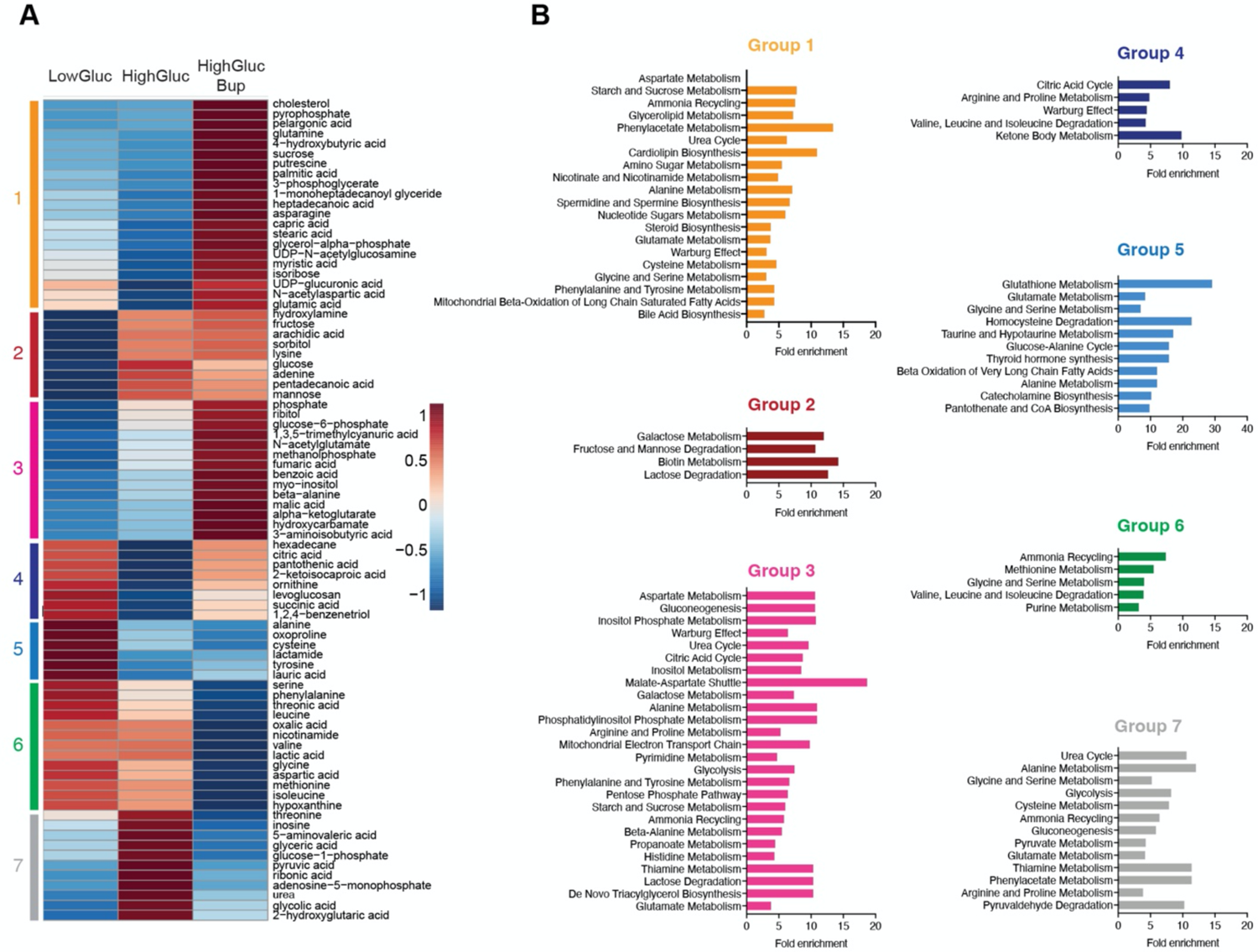
Unbiased primary metabolite profiling of glucose- and BP-treated SCs identifies treatment-specific pathway modulations. **A)** Average level of the primary metabolites quantified in three biological replicates. **B)** Pathway analysis of the metabolite clusters using Metaboanalyst (Xia et al., 2009). Identified primary metabolites of differentially treated SCs fall into seven categories based on their abundance pattern. Groups 4 and 7 show metabolites that their enrichment or depletion under hyperglycemia is reversed upon BP treatment.

**Figure S11:**
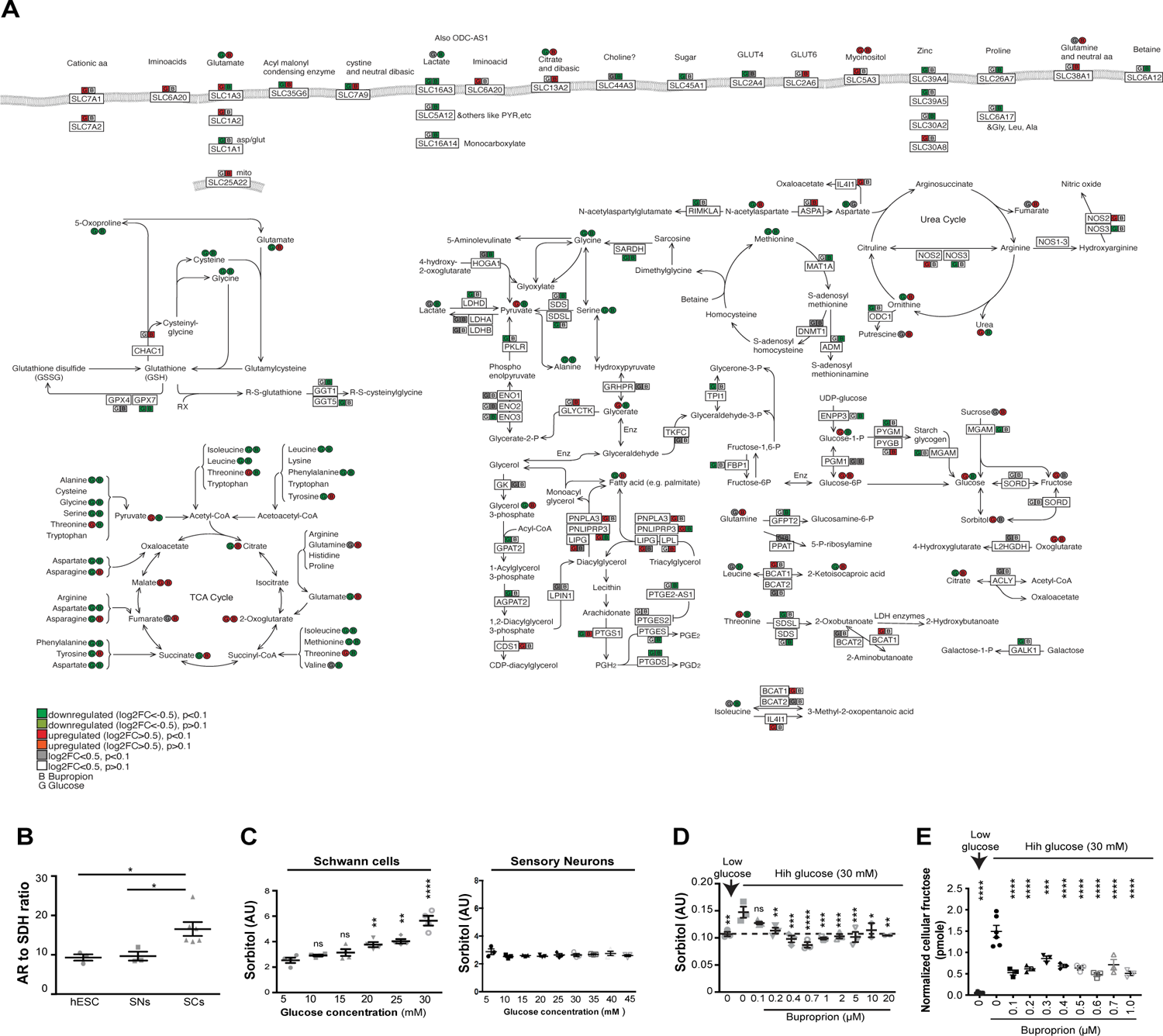
Altered metabolic pathways in SCs treated with high glucose and BP. **A)** RNA sequencing and metabolomics data were integrated to draw this schematic according to metabolic pathways in KEGG database. **B-E**) Polyol pathway is affected by glucose and BP treatments. **B**) qPCR analysis of Aldose Reductase (AR) to Sorbitol Dehydrogenase (SDH) ratio in undifferentiated cells, SCs and sensory neurons. p-values are: p-value: * p<0.05; ** p<0.01. **C**) Intracellular sorbitol measurements in hPSC-derived SCs and sensory neurons in response to exposure to different glucose concentration. p-values are: p-value: * p<0.05; ** p<0.01. **D**) Dose response analysis of BP for effects on sorbitol levels. **E**) Dose response analysis of BP for effects on fructose levels. ns, not significant, p-values are: * p<0.05; ** p<0.01; *** p<0.001; **** p<0.0001.

**Figure S12:**
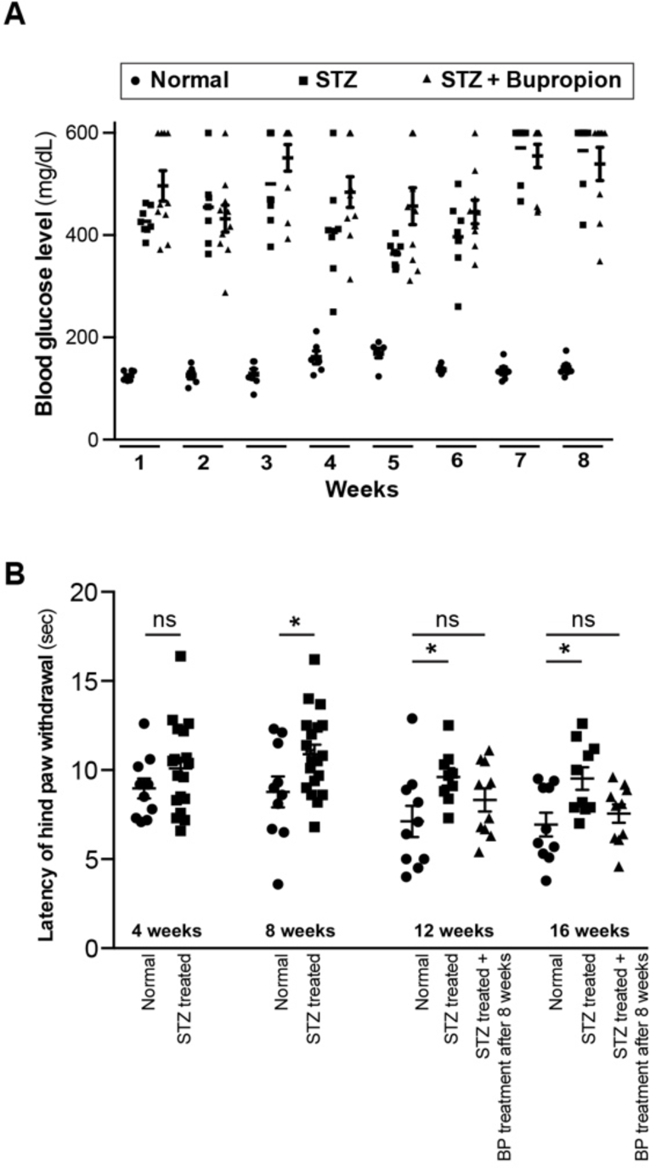
BP treatment doesn’t affect STZ-induced hyperglycemia but reverses nerve damage in mice. **A)** Blood glucose levels in normal mice and mice treated with STZ and Bupropion. **B)** Thermal sensitivity test measuring the latency of hind paw withdrawal in normal mice, mice treated with STZ only and mice treated with STZ then with BP starting at 8 weeks post STZ treatment. p-values: * p<0.05.

## SUPPLEMENTARY TABLES

**Table S1:** List of gene categories used for characterization of SC lineages

**Table S2:** List of SC lineage markers generated from primary mouse datasets

**Table S3:** List of markers for myelinating SCs.

**Table S4:** List of normalized z-scores from the high-throughput screen

**Table S5:** List of all predicted targets for hit compounds

**Table S6:** List of primary antibodies and working dilutions

**Table S7:** Clustering parameters for single cell RNA sequencing datasets

## METHODS

### Culture of human pluripotent stem cells (hPSCs)

hPSC line H9 (WA-09) and derivatives (*SOX10::GFP; SYN::ChR2-YFP; SYN::YFP;PHOX2B:GFP; EF1::RFP EDNRB-/-*) were maintained on mouse embryonic fibroblasts (MEF, Global Stem, Rockville, MD) in KSR (Life Technologies, 10828-028) containing hPSC medium as described previously (Chambers et al., 2009) or were plated on geltrex™-coated plates and maintained in chemically-defined medium (E8) as described previously (Barber et al., 2019). Cells were subjected to mycoplasma testing at monthly intervals and STR profiled to confirm cell identity at the initiation of the study.

### Neural crest induction

Differentiations of hPSCs towards NC were carried out following previously established methods using the knockout serum (KSR) medium or chemically defined Essential 6 (E6) medium (Fattahi et al., 2016; Tchieu et al., 2017). Briefly, when the monolayer culture of hPSCs reached about 70% confluency, neural crest induction protocol was initiated (D0) by aspirating the maintenance medium (E8) and replacing it with neural crest induction medium A [BMP4 (1 ng ml^-1^), SB431542 (10 μM), and CHIR 99021 (600 nM) in Essential 6 medium]. Subsequently, on D2 neural crest induction medium B [SB431542 (10 μM) and CHIR 99021 (1.5 μM) in Essential 6 medium] was fed to the cultures until D12. Next, developing precursors are formed during D12–D30 to facilitate the selection for glial progenitor lineages. In doing so, we removed medium B on D12 and detached the NC monolayers using accutase (30 min, 37 °C, 5% CO_2_). After centrifuging the samples at 290 x g for 1 min, we re-suspended the cells in NC-C medium [FGF2 (10 ng ml^-1^), CHIR 99021 (3 μM), N2 supplement (10 μl ml^-1^), B27 supplement (20 μl ml^-1^), glutagro (10 μl ml^-1^), and MEM NEAAs (10 μl ml^-1^) in neurobasal medium] and transferred them to ultra-low-attachment plates to form free-floating 3D developing precursors. Two days later, when the free-floating developing precursors could be observed, we gently gathered them in the center of each well using a swirling motion. Then, the old media was carefully aspirated from the circumference of each well without removing developing precursors. After addition of the fresh NC-C medium, the cultures were incubated for 48 hours (37 °C and 5% CO_2_) prior to passaging using accutase. Similarly, cultures were fed with fresh medium every other day and passaged every four days until D30.

### Induction and expansion of Schwann cells from hPSCs

At day 1, NC cells were aggregated into 3D spheroids (5 million cells/well) in Ultra Low Attachment 6-well culture plates (Fisher Scientific, 3471) and cultured in Neurobasal (NB) medium supplemented with L-Glutamine (Gibco, 25030-164), N2 (Stem Cell Technologies, 07156) and B27 (Life Technologies, 17504044) containing CHIR (3 µM, Tocris Bioscience, 4423) and FGF2 (10 ng/ml, R&D Systems, 233-FB-001MG/CF) and NRG1 (10 ng/ml, R&D 378-SM-025). After 14 days of suspension culture, the spheroids were plated on Poly Ornithine/Laminin/ Fibronectin (PO/LM/FN) coated dishes (prepared as described previously (17)) in Neurobasal (NB) medium supplemented with L-Glutamine (Gibco, 25030-164), N2 (Stem Cell Technologies, 07156) and B27 (Life Technologies, 17504044) containing NRG1 (20 ng/ml, R&D 378-SM-025), FGF2 (10 ng/ml, R&D Systems, 233-FB-001MG/CF) and cAMP (100 mM, Sigma, D0260). The SC precursors migrate out of the plated spheroids and differentiate into SCs within 10 days. For long-term expansion, cells were cultured in Schwann cell medium (Sciencell, 1701) on PO/LM/FN coated dishes. Cells were fixed for immunostaining or harvested for gene expression analysis at Day 25, Day 35, Day 50 and Day 100 of differentiation.

### FACS and Immunofluorescence (IF) analysis

For IF, the cells were fixed with 4% paraformaldehyde (PFA, Affymetrix-USB, 19943) for 20 minutes, then blocked and permeabilized using 1% Bovine Serum Albumin (BSA, Thermo Scientific, 23209) and 0.3% triton X-100 (Sigma, T8787). The cells were then incubated in primary antibody solutions overnight at 4**°**C (Celsius) and stained with fluorophore conjugated secondary antibodies at RT for 1 hour, the stained cells were then incubated with DAPI (1 ng/ml, Sigma, D9542-5MG) and washed several times before imaging. For Flow Cytometry analysis, the cells are dissociated with Accutase (Innovative Cell Technologies, AT104) and fixed and permeabilized using BD Cytofix/Cytoperm (BD Bioscience, 554722) solution, then washed, blocked and permeabilized using BD Perm/Wash buffer (BD Bioscience, 554723) according to the manufacturer’s instructions. The cells are then stained with primary (overnight at 4) and secondary (30 min at room temperature) antibodies and analyzed using a flow Cytometer (FlowJo software). A list of primary antibodies and dilutions is provided in Table S6.

### Single cell RNA-sequencing (scRNA-seq) data analysis ScRNA-seq Processing

FASTq files were aligned using the 10X Genomics CellRanger 6.0.0 pipeline (Zheng et al., 2017) to the human GRCh38 reference transcriptome to generate gene-expression counts matrices using the “include introns” option.

### Quality Control and Cell Filtration

Datasets were analyzed in R v4.1.0 with Seurat v4 (Hao et al., 2021). The number of reads mapped to mitochondrial and ribosomal transcripts per cell were derived using the “PercentageFeatureSet” function. We identified cells of poor quality and subsequently removed them independently for each dataset based on the number of unique features captured per cell, the number of unique molecular identifiers (UMI) captured per cell and the percentage of mitochondrial gene transcripts per cell. Datasets were filtered based on the following quality control metrics: nFeatures>200, nFeatures<7000, nCounts< 40000 and percent mitochondrial reads < 20%.

### Dimensionality Reduction, Clustering and Annotation

Transcript count matrices were log normalized applying a scaling factor of 10,000 with 2,000 variable features identified using the “vst” method. Cell cycle phase distribution was predicted using the “CellCycleScoring” function with Seurat’s S and G2M features available in “cc.genes”. The 2000 most variable feature set was scaled and centered, and the following data variables were regressed out: nFeatures, nCounts, mitochondrial gene percentage, ribosomal gene percentage, S score and G2M score. Principal Components Analysis (PCA) was run using default settings. PCA reduction was used to perform Uniform Manifold Approximation and Projection (UMAP) dimensionality reduction. The shared nearest neighbors (SNN) graph was computed using default settings followed by cell clustering achieved using the default Louvain algorithm. Quality control metrics were visualized per cluster to identify and remove clusters of low-quality cells (see Quality Control and Cell Filtration). The above pipeline was performed again on each dataset after the removal of low-quality cell clusters. The number of principal components used for UMAP reduction and SNN calculation was determined by principal component standard deviation unique for each dataset together with resolution used for clustering of each dataset can be found in Table S7. Cluster markers were derived using the Wilcoxon Rank Sum test. Cluster annotation was based on the expression of known cell type marker genes. Following cell type annotation, gene dropout values were imputed using adaptively-thresholded low rank approximation (ALRA) (Linderman et al., 2018). The rank-k approximation was automatically chosen for each dataset with default values selected for all other parameters. The imputed gene expression is depicted in all plots and used as default in all downstream analyses unless otherwise specified

### Analysis of Published Datasets

Primary tissue derived Schwann cell type markers for Schwann cell precursor, myelinating and non-myelinating Schwann cells were obtained from Tasdemir-Yilmaz et.al. (Tasdemir-Yilmaz et al., 2021) using interactive webpage Pagoda 2 by performing differential feature expression analysis of cell type cluster of interest against the entirety of dataset cells. Differentially expressed (DE) genes were sorted by Z-score and converted to human gene names with “biomaRt” (Durinck et al., 2009) package using human and mouse genome databases available at ensembl.

### Gene Group Expression Characterization

Gene lists were compiled for genes belonging to transcription factor, surface marker, cell adhesion, neurotransmitter receptor and neurotrophic factor functional groups from Molecular Signatures Database (MSigDB) (Liberzon et al., 2011). For each dataset, the gene lists were filtered to remove low abundance genes (detected in less than 25% of cells of each cell type cluster). Genes were then determined to be exclusively expressed by a cluster if greater than 25% of cells within that cluster only expressed the gene. To further selectively filter transcription factor and surface marker gene sets we derived genes shared by transcription factor and surface marker gene sets and cell type specific differentially expressed (DE) gene lists.

### Cell Type Transcriptional Signature Scoring

To find transcriptionally similar cell populations between two datasets, first the differentially expressed (DE) genes of the reference dataset were calculated from the imputed gene counts with the “FindAllMarkers” function using the Wilcoxon Rank Sum test and only genes with a fold change (FC) above 0.25 were returned. The reference DE gene lists were filtered to remove genes not present in the query dataset. Then for each cell cluster in the reference dataset, a transcriptional signature gene list was made from the top 100 DE genes filtered for p value below 0.05 and sorted by decreasing fold change (FC). The query dataset is then scored for the transcriptional signature gene lists of each reference dataset cell cluster using the “AddModuleScore” function based on the query dataset’s imputed feature counts.

### Myelinating Schwann Cell Identity Specification

Myelinating Schwann cell (mySC) specific marker gene set was curated by combining published dataset derived markers (Jessen and Mirsky, 2005) and canonical myelination associated markers for a total of 21 marker genes (Table S3). LP and HP specific mature Schwann cell clusters were subset from entire LP and HP datasets and scored for mySC gene set using the “AddModuleScore” function. Cells with positive mySC score were isolated and identified as mySC in further analyses.

### Gene Ontology Analysis

Cell type of interest specific DE genes with positive fold change (FC) were calculated from the imputed gene counts with the “FindAllMarkers” function. Each gene set was filtered to include genes with p value<0.05 and sorted by decreasing fold change. Where possible up to 250 genes from each cell type specific dataset were used in gene functional profiling analysis by using g:Profiler (Raudvere et al., 2019) online tool and selecting pathways from GO biological process, KEGG and Reactome databases. Term enrichment was ranked by decreasing value of negative log10 transformed p values.

### Surface marker screening

Screening for specific surface antigens was performed using BD Lyoplate library**^®^** (BD, 560747) on hPSC-SCs at day 80 of differentiation. Cells were plated in 96 well plates (10,000 cells/well) and stained with primary and secondary antibodies according to manufacturer’s instructions. The stained wells were fixed for total plate imaging and quantification. The percentage of double positive cells out of total GFAP was quantified for each antibody. Top hits (>60% double positive) were validated further using flow cytometry.

### Gene expression analysis

For RNA sequencing, total RNA was extracted using RNeasy RNA purification kit (Qiagen, 74106). For qRT-PCR assay, total RNA samples were reverse transcribed to cDNA using Superscript II Reverse Transcriptase (Life Technologies, 18064-014). qRT-PCR reactions were set up using QuantiTect SYBR Green PCR mix (Qiagen, 204148). Each data point represents three independent biological replicates. RNA-seq reads were mapped to the human reference genome (hg19) using TopHat v2.0. TopHat was run with default parameters with exception to the coverage search. Alignments were then quantified using HTSeq and differential gene expression was calculated using DESeq normalized to the cranial neural crest sample.

### Viability assay

To monitor the viability of SCs, cells were assayed for LDH activity using CytoTox 96 cytotoxicity assay kit (Promega, G1780). Briefly, the cells are plated in 96 well plates at 30,000 cells/cm^2^. The supernatant and the cell lysate is harvested 24 hours later and assayed for LDH activity using a plate reader (490 nm absorbance). Cytotoxicity is calculated by dividing the LDH signal of the supernatant by total LDH signal (from lysate plus supernatant). The cells were cultured in Schwann cell medium (Sciencell, 1701) on PO/LM/FN coated dishes during the assay.

### Calcium imaging

MN-only cultures and MN-SC co-cultures were subjected to calcium imaging at days 40 and 70 post-co-culture as previously described (Barreto-Chang and Dolmetsch, 2009). Briefly, The cells were loaded with 2 μmol/L Fluo-4 AM dissolved in 1:1 (v/v) amount of 20% Pluronic®-F127 and DMSO with stock concentration of 1 mmol/L for 45 min at RT in Tyrode solution consisting of (mmol/L): 140 NaCl, 5.4 KCl, 1 MgCl2, 1.8 CaCl2, 10 glucose and 10 HEPES at pH 7.4. For activation, cells were spiked with a solution containing glutamate (50 mM) or KCl (300 mM). Time lapse images were acquired using an Axiovert Inverted Microscope (Zeiss) on a heated stage. Ratiometric analysis was performed using Metamorph Software (Molecular Devices).

### Transplantation of hPSC-SCs in rat sciatic nerves and histological assessment

All procedures were performed following NIH guidelines, and were approved by the local Institutional Animal Care and Use Committee (IACUC). Rats were placed under isoflurane gas anesthesia and both sciatic nerves were exposed below the sciatic notch and crushed using Dumont #5 forceps for 30 seconds twice in the same location. Immediately afterwards, a cell suspension of 3×104 hPSCs/μl Schwann cells were transplanted via injection of ∼3-4 μl at proximal and distal locations to the crush site with a glass micropipette. Survival times ranged from 2 to 8 weeks. For immunohistochemistry, tissue was fixed through intracardiac perfusion of 4% PFA in 0.1 M PBS. Sciatic nerves were dissected from rats at 2, 3, 4, 8 weeks after crush lesion and transplantation. After dissection, sciatic nerves where prepared by placing them in 30% sucrose in 0.1 M PBS overnight and embedding them in OCT blocks for cryosectioning, or, by removing the perineurium and teasing them in cold 0.1 M Phosphate Buffer (pH 7.4). Some nerves were teased after perfusion and immunostained to examine individual axons. Regenerated axons distal to the crush site were analyzed.

### Metabolite measurements

High glucose, low glucose and drug treated SCs and sensory neurons were subjected to biochemical assays for Sorbitol (Abcam, ab118968), glucose (Abcam, ab65333), Pyruvate (Abcam, ab65342) and 2DG uptake (Abcam, ab136955). The measurements were carried out according to the manufacturer’s instructions. Data were normalized according to cell numbers and averaged across 3-6 biological replicates.

### High-throughput screening assay for drugs reversing glucose-mediated SC cytotoxicity

The chemical compound screening was performed using the Prestwick Chemical Library**^®^**. RFP-labeled hPSC-SCs were plated in 384 well plates (1,000 cells/well) and treated with 30 mM glucose immediately before addition of the compounds. The compounds were added at 1 µM concentration. After 72 hours, the plates were treated with DAPI for ten minutes, washed twice and fixed for total plate imaging. The number of viable cells was quantified for each well by counting the number of DAPI negative, RFP positive cells. For validation of the selected hit compound (Bupropion HCL, Sigma, B102), the cells were treated with various concentrations of the compound for dose response analysis. The highest non-toxic dose (0.7 µM; based on sorbitol reduction and viability) was used for follow-up experiments.

### Drug target prediction

Z-scores for primary hit compounds were calculated as Z= (x-µ)/σ. X is the number of viable cells. µ is the mean number of viable cells and σ is the standard deviation for all compounds and DMSO controls. The normalized *z*-score values reported for all the compounds were first transformed to N(0,1) using the bestNormalize package (v1.4.0) in R (v3.5.1). The treatments with transformed *z*-scores greater than 2 were selected, which resulted in 16 hit compounds.

To identify the proteins that are most likely targeted by the SC protecting drugs against glucotoxicity, we employed two independent tests. First, we calculated combined z-score in which we integrated normalized z-score from all of the treatments associated with a particular protein. Second, we performed a Fisher’s exact test to see if a target protein is enriched among the targets of treatments with positive z-scores. We report the correlation between the *p-*-values calculated by these two independent tests. For every compound, possible target proteins were identified as above. Weighted combined z-scores were then calculated for each protein by combining normalized *z*-scores across all treatments (Zaykin, 2011). The *p*-values were then calculated based on the combined z-scores and adjusted using p.adjust (method=FDR). As an orthogonal approach, for each protein, we recorded the number of treatments with positive normalized *z*-scores as well as the total number of compounds predicted to target that protein. Using the sum of counts for all other proteins and drugs, we performed a Fisher’s exact test to evaluate the degree to which positive *z*-scores were enrichment among the treatments likely to affect a protein of interest. As expected, the two *p*-values, i.e. combined *z*-score and Fisher’s, are generally correlated.

### Protein-protein interaction network construction

Protein-protein interaction network analysis was performed using the Search Tool for the Retrieval of Interacting Genes (STRING) database. The minimum required interaction score was set to 0.4 corresponding to medium confidence. The edge thickness indicates the degree of data support from the following active interaction sources: textmining, experiments, databases, co-expression, neighborhood, gene fusion and co-occurrence.

### iPAGE pathway enrichment analysis

The iPAGE algorithm was used for gene set and pathway enrichment analysis (Goodarzi et al., 2009). iPAGE first quantize continuous input data into equally populated bins and then, it calculates the Mutual Information (MI) between a vector of values for genes within each cluster bin and a binary vector of gene set memberships. The significance of the calculated MI values is then assessed through a randomization-based statistical test. Finally, it uses hypergeometric distribution to determine the level with which the significantly informative pathways are overrepresented (red) or underrepresented (blue) in each cluster bin. The resulting p-values asses to draw heatmap visualization, in which rows represent significant pathways and columns correspond to cluster bins. We first ordered target genes identified from FDA approved drug library screening with positive z-score (see drug target prediction above) based on their combined z-score from left to right and followed by dividing them into seven equally populated bins. We then assessed the enrichment (red boxes) and depletion (blue boxes) of various gene sets across the spectrum (MSigDB v6.0). Gene sets provided were from the Molecular Signatures Database (MSigDB v6.0) database (Liberzon et al., 2015). Here, we report enriched gene sets from cluster 2 (C2, curated gene sets), and cluster 5 (C5, ontology gene sets).

### NF-kB staining

Image analysis was performed using FIJI (Schindelin et al., 2012). Watershedding was applied to separate nuclei that were grouped together. Nuclear ROIs were overlayed onto the NF-κB image to calculate nuclear co-localization. To analyze NF-κB in each condition, several measures were taken. Because the cytoplasm could not be isolated to represent the cell to which it belongs and because the average of the mean intensity of the cytoplasmic ROIs would not account for the distribution of intensities per ROI area, the integrated densities of the cytoplasmic ROIs were summed together and divided by the sum of the cytoplasmic ROIs’ area to give the mean intensity of NF-κB across all cytoplasms in each image. The mean intensities of each nuclei were divided by the mean intensity of the cytoplasm to give a ratio of nuclear to cytoplasmic NF-κB expression.

### Metabolomics

hPSC-Scs were treated with 5mM and 30mM glucose for 72 hours and harvested for metabolomics analysis. Frozen total cell pellets from at least three biological repeats were submitted to the West Coast Metabolomics Center at the University of California, Davis that used Agilent 6890 gas chromatographer and Pegasus III TOF mass spectrometer for an untargeted primary metabolomics analysis. Metaboanalyst (Xia et al., 2009) was used to generate the heat maps and perform path analysis.

### Fructose, lactate and sorbitol measurements

Cellular levels of fructose, lactate and sorbitol were measured by colorimetric assays following the instructions provided by the manufacturers. The following kits were used: fructose (Abcam, ab83380), lactate (Abcam, ab65331) and sorbitol (Abcam, ab118968)

### CRISPR knockout of PTGER4

Schwann PTGER4 ribonucleoprotein (RNP) complex was assembled by mixing 180 pmol of multiguide sgRNA (Synthego, USA) and 20 pmol of Cas9 2NLS (Berkeley QB3) in Lonza electroporation buffer P3 (Lonza, Switzerland) per reaction. hPSC-derived Schwann cells were detached using trypsin and washed 2 x with PBS. 250,000 cells per reaction were resuspended in Lonza electroporation buffer P3 immediately before electroporation. Cells were mixed with the RNPs and were electroporated using a Lonza 4D 96 well electroporation system with pulse code DS-137. Ten minutes post nucleofection, cells were then diluted in warm medium and were plated. Medium was changed on the following day, cells were passed as needed until the day of the assay. For quantification, images were analyzed with NIH ImageJ software by measuring the fluorescence intensity of individual cells by manual region of interest selection.

### Drug treatment of diabetic mice

All procedures were performed following NIH guidelines, and were approved by the local Institutional Animal Care and Use Committee (IACUC). 3-8 weeks old male C57BL6 mice were treated with one dose IP injection of STZ (180 mg/kg, sigma, 85882) to induce pancreatic beta cell death. Blood glucose levels were measured using a standard Glucometer (Freestyle Lite) in weekly intervals starting at one week post-treatment, by drawing a drop of blood from tail tip. BP treatment was initiated one week post-STZ treatment. BP was mixed with standard chow at 1.63 mg/g of chow to be administered orally at ∼300 mg/kg daily. The dose was calculated based on average daily food intake (5.5 g/day) and initial body weight (30 g).

### Mouse thermal sensitivity test

Thermal nociception was assessed using the hot plate test. The hot plate (Ugo Basile 35100) consisted of a metal surface (55 °C) with a transparent Plexiglas cylinder to contain the mouse. The subject is placed upon a constant temperature hot plate and the latency necessary to demonstrate discomfort, assessed by either licking/shaking the hind paw or jumping, is determined. Typical baseline latencies are 5-10 seconds, with maximal latencies of 30 seconds. Any animal that does not demonstrate discomfort behavior will be removed after 30 seconds, the maximal latency, to avoid tissue damage.

### Statistical analysis

Data are presented as mean ± SEM and were derived from at least 3 independent experiments. Data on replicates (n) is shown in figures. Statistical analysis was performed using the Student t-test (comparing 2 groups) or ANOVA with Dunnett test (comparing multiple groups against control). Distribution of the raw data approximated normal distribution (Kolmogorov Smirnov normality test) for data with sufficient number of replicates to test for normality.

